# Secretome of Human Umbilical cord mesenchymal stem cells exerts protective impacts on the blood-brain barrier against alpha-synuclein aggregates using an *in vitro* model

**DOI:** 10.1101/2023.10.21.562544

**Authors:** Kimia Marzookian, Farhang Aliakbari, Hamdam Hourfar, farzaneh sabouni, Daniel E. Otzen, Dina Morshedi

## Abstract

The blood-brain barrier (BBB) is a highly developed endothelial microvessel network extended to almost all parts of the central nervous system (CNS) that tightly seals cell-to-cell contacts and plays a critical role in maintaining CNS homeostasis. It also protects neurons from factors present in systemic circulation and prevents pathogens from entering the brain. Conversely, BBB disruption can initiate multiple pathways of nerve damage. BBB injury contributes significantly to various neurodegenerative diseases, including Parkinson’s disease (PD). PD is also characterized by aggregation of the protein alpha-synuclein (αSN) to form intracellular inclusions. Recent studies have shown that due to their active secretions, mesenchymal stem cells (MSCs) can effectively relieve the severity of many neurological diseases. However, the impact of MSCs on BBB remains largely unclear. Here, we investigated the effect of Secretome extracted from MSCs on BBB when treated with toxic αSN-aggregates (αSN-AGs). For this purpose, MSCs were first isolated from Umbilical cord tissue (UC-MSC), and their secretome was collected. Then, the impact of the secretome on the cytotoxicity and inflammatory effects of αSN-AGs was examined on hCMEC/D3 cells using in vitro BBB models produced by mono- and co-culture systems. We explored the effects of αSN-AGs in the presence of UC-MSC secretome on permeability, TEER value, and cytokine/chemokine release. We found that the Secretome of UC-MSCs exerts protective effects by inhibiting the toxic effects of αSN-AGs on the BBB. These results strongly support the potential of UC-MSCs secretome for cell-free PD therapy. We also present an improved method for isolation of MSCs from umbilical cord tissue, which we hope will facilitate further studies on the use of these cells.

**Graphical Abstract:** 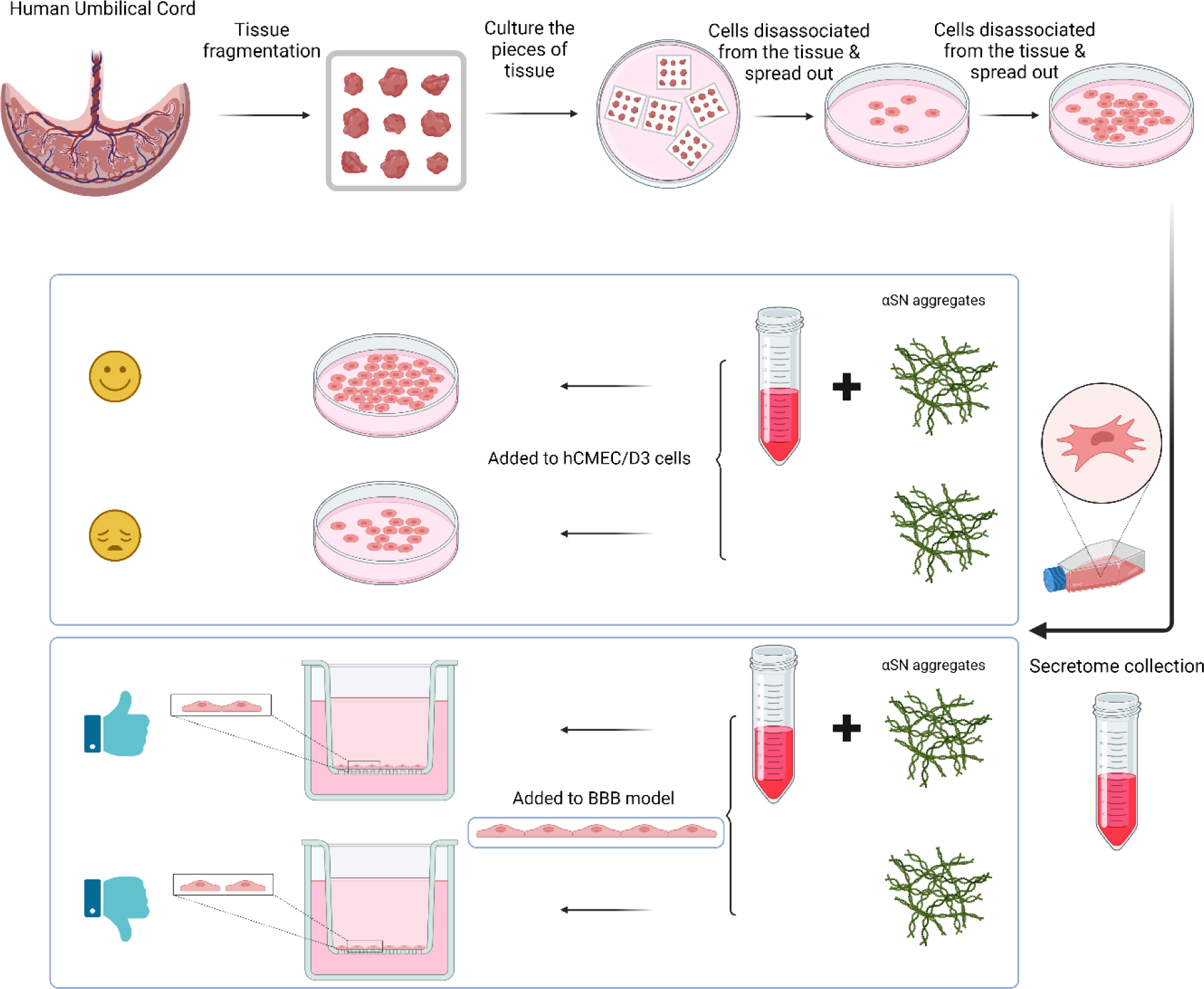

## 1. Introduction

Parkinson’s disease (PD) is the second most prevalent progressive neurodegenerative disease worldwide, characterized by the loss of dopaminergic neurons in the substantia nigra pars compacta (SNpc) and depletion of dopamine concentration in the striatum (Fearnley and Lees, 1991). The worldwide prevalence of PD has increased from two and a half million people in 1990 to more than 6 million people in 2016 (Dorsey *et al*., 2018), and is estimated to reach 13 million people by 2040 (Jankovic and Tan, 2020). The leading cause of PD is related to the aggregation of the protein alpha-synuclein (αSN). Throughout the progression of the disease, αSN converts into various aggregated conformers, which predominantly present as protein plaques-like β-sheet structures called Lewy bodies (LB) and Lewy neurites (LN) in the neurons in PD patients (Spillantini *et al*., 1997). However, these phenomena are not confined to the SNpc. αSN aggregates (αSN-AGs) are expanded in various parts of the brain, including the brainstem, anterior olfactory structures, and medulla, before being detected in the SNpc. LB and LN have also been identified in the brain cortex in the late stages of the disease (Braak et al., 2006). In addition, αSN and its oligomeric species have been detected in PD patients’ cerebrospinal fluid (CSF) and blood, suggesting that it is able to cross the BBB (El-Agnaf *et al*., 2003)(Tokuda *et al*., 2010).

The BBB comprises microvascular endothelial cells (BMECs) connected by tight junctions with transmembrane and integral membrane proteins, forming an impermeable structure in brain blood vessels. This barrier, along with neurons, pericytes, astrocytes, microglia, and oligodendrocytes, creates the neurovascular unit (NVU). Together, they preserve the nervous tissue’s homeostasis. (Lauschke, Frederiksen and Hall, 2017). There is strong evidence that in neurodegenerative diseases, including Parkinson’s, Alzheimer’s, and multiple sclerosis, BBB dysfunction plays a critical role in the pathogenesis and spread of the disease (Stolp and Dziegielewska, 2009)(Desai *et al*., 2007)

Toxicity due to αSN-AGs in the brain and the blood of Parkinson’s patients has a direct relation with neurodegeneration in which neuroinflammation and dysfunction of the BBB may be involved (Al-Bachari *et al*., 2017)(Wu *et al*., 2021). To better understand the mechanisms involved in αSN pathology on the BBB and dopaminergic neurons, Pediaditakis et al used organ-on-a-chip technology to show that αSN-AGs causes mitochondrial dysfunction and apoptosis in neuronal cells, increase neuroinflammation through the upsurge of pro-inflammatory factors and BBB dysfunction leading to an increase in its permeability. They also found that the dysfunction of the BBB was due to the effect of the αSN-AGs on the expression of junction proteins such as zonula occludens and claudin family in the brain capillary endothelial cells and increased expression of pro-inflammatory cytokines such as TNF-α and IL-6 (Pediaditakis *et al*., 2021).

The vast majority of PD cases (∼90%) is non-hereditary (idiopathic) while only 10% are associated with mutations in one of the 20 known genes associated with this disease. The underlying processes usually start several years before the symptoms are observed (Chen *et al*., 2022). Although the exact mechanism of PD and its propagation in the brain have not yet been determined, Braak and have proposed that αSN-AGs form outside the brain, *e.g.* the olfactory or enteric nerve cell plexuses and only the final stages of the disease in the brain lead to visible clinical symptoms (Braak *et al*., 2003). This theory was later expanded to a “dual hit” hypothesis in which αSN-AG effects are augmented by a neurotropic pathogen, probably viral, which enters the brain via nasal or gastric routes at places where the vascular barrier has been breached (Hawkes, Del Tredici and Braak, 2007)

Also, there is evidence that environmental factors increase the risk of PD, including toxins such as pesticides, air pollutants, metals, and smoking. According to the dual hit hypothesis, these factors may trigger inflammation and spread via gut-to-brain pathways to initiate or exacerbate the abnormal aggregation of α-synuclein (Hawkes, Del Tredici and Braak, 2009)(Chen *et al*., 2022).

We have recently shown that αSN-AGs species disrupt the function and integrity of BBB in an in vitro model. We attribute this to aberrant angiogenesis through induction of TNF-α expression in PD patient-derived brain endothelial cells (Hourfar *et al*., 2023). Given the significance of the BBB in PD progression and the impact of αSN-AGs on BBB integrity, it is essential to develop novel approaches to minimize BBB damage and enhance its protection.

Mesenchymal stem cells, with their unique characteristics, can be a versatile therapeutic strategy against PD symptoms (Zhou *et al*., 2022)(Malekpour *et al*., 2022). Mesenchymal stem cells (MSCs) are pluripotent stromal cells that persist to a small extent in some mature tissues and have a fibroblast-like morphology, with the ability to self-renew and differentiate into different tissues (Uccelli, Moretta and Pistoia, 2008) MSCs can trans/differentiate to neural precursors or mature neurons and promote neuroprotection and neurogenesis. MSCs are characterized by the expression of specific surface markers, including CD105, CD73, and CD90, and the lack of expression of CD45, CD34, CD14, CD11b, and CD19. They are mainly collected from adipose tissue, umbilical cord, placenta, and bone marrow for clinical and research purposes (Young *et al*., 1995)(Berebichez-Fridman and Montero-Olvera, 2018).

MSCs transplantation has positively affected BBB integrity and dopaminergic neuron regeneration in PD animals (Chao, He and Tay, 2009). MSC transplantation can recover degenerative nerves by migrating into damaged areas and contributing to neuron regeneration through paracrine activity and differentiation into neural-like cells (Venkatesh and Sen, 2017). However, this method is still not considered a standard clinical treatment due to some shortcomings, including loss of activity in long-term maintenance, lack of standard instructions for expanding cells in the body, lack of knowledge about the correct injection method, optimal injection dose, and administration frequency. Furthermore, MSC transplantation is compromised by the potential risk of side effects like tumorigenesis. Consequently, medical communities are shifting their focus toward cell-free treatment methods (Kumar *et al*., 2019) (Meiliana, Dewi and Wijaya, 2019). This shift in focus is encouraged by the observation that MSCs exert most of their properties (*e.g.* anti-apoptotic, anti-inflammatory and immune system-modulating properties as well as effects on implantation and migration) through their secreted substances, collectively called the secretome, which include growth factors, extracellular vesicles, cytokines, and chemokines (Teixeira *et al*., 2013)(Madrigal, Rao and Riordan, 2014).

Currently, common clinical treatments for PD involve dopamine analogs and deep brain stimulation, which only reduce the symptoms of the disease but do not stop its progress (Sethi, 2010)(Muthuraman *et al*., 2018). PD animal model studies demonstrated that the Secretome of MSCs effectively mitigated both apoptosis and damage to dopaminergic neurons. Furthermore, the administration of MSCs’ Secretome successfully ameliorated motor deficits in a mouse model of PD (Mendes-Pinheiro *et al*., 2019)(Chen *et al*., 2020).

The growing understanding of the simultaneous presence of αSN in the nervous system and surrounding tissues and its propensity to form toxic aggregates has underscored the urgency to investigate the communication between αSN and the BBB closely. This intricate connection demands careful attention to fully comprehend the implications and potential therapeutic opportunities for managing αSN-related disorders (Sui *et al*., 2014).

αSN monomers and oligomers are able to pass through the BBB through a clathrin-dependent pathway in the direction of the luminal-abluminal pathway (Alam *et al*., 2022). Given this targeting of the BBB by αSN-AGs, we hypothesize that avoiding or reducing this damage may prevent the PD from spreading throughout the brain.

In this study, we investigate the protective effects of the hUC-MSCs-derived secretome on an in vitro model of the BBB treated with toxic αSN-AGs through the laminal direction, which, to the best of our knowledge, has not been reported before.

## 2. Materials and Methods

### 2.1. Reagents and chemicals

3-(4,5-dimethylthiazol-2-yl)-2,5-diphenyl tetrazolium bromide (MTT), Griess reagents, gelatin, Poly-L-Lysine, Bovine type I collagen, hydrocortisone, 4’,6-Diamidino-2-phenylindole (DAPI), Thioflavin T (ThT) purchased from Sigma-Aldrich (USA). The BCA Protein Assay Kit (Thermo Fisher, USA). Roswell Park Memorial Institute (RPMI) 1640 medium, DMEM/F-12 (Dulbecco’s Modified Eagle Medium/Nutrient Mixture F-12), and DMEM high glucose and antibiotics were from GibcoBRL. All salts and organic solvents were from Merck (Darmstadt, Germany). Fetal bovine serum (FBS) was from Biosera (Tehran, Iran). Transwell 24-well tissue culture inserts (0.4 μm pore size, polyester membrane) were from Plastics & Membrane Technology (SABEU, Germany). Human TNF-alpha, IL-10, IL-8, CCL2 Quantikine ELISA Kit (R&D systems quality control, USA).

### 2.2. Methods

#### 2.2.1. MSCs isolation and characterization and collection of their Secretome

##### 2.2.1.1 Isolation of MSCs from human umbilical cord

The umbilical cord connected to the placenta junction was obtained from a healthy mother after cesarean delivery. Consent was obtained from the parents and hospital under the standards guidelines of the National Institute of Genetic Engineering and Biotechnology ethics committee (ethics number: IR. NIGEB.EC.1399.4.9.C). Tissues were transferred to the laboratory at 4 °C in a sterile container containing PBS with antibiotics (ciprofloxacin 10 μg/mL, gentamicin 50 μg/mL, amphotericin B 10 μg/mL). The cells were extracted up to 4 h after acquiring the umbilical cord. Under a laminar hood, the tissue was washed several times with sterile PBS containing antibiotics to remove blood clots; then, the placenta junction was separated from the umbilical cord. The umbilical cord was then divided into 1-2 cm pieces, and after removing the blood vessels, Wharton jelly, umbilical cord lining, and placenta junction sections were cut into 1-2 mm pieces and cultured by an explant method in the bottom of 10 cm^2^ plates in DMED-F12 culture medium with 15% FBS and 100 U/mL penicillin, and 100 μg/mL streptomycin. Finally, the cultures were incubated in a humidified incubator for 12-14 days (5% CO_2_ and 90% moisture). In Fig. S1, the detail of the modified explant method is demonstrated.

##### 2.2.1.2. Optimizing Culture Conditions for Improved Extraction of MSCs from the Umbilical Cord

Since explant culture is superior to enzymatic method in terms of yield of isolation and viability of MSCs, we chose this method for MSC isolation (Yoon *et al*., 2013)(Hendijani, 2017).

In this method, it is necessary to attach tissue pieces to a culture dish in order to migrate the MSCs from the pieces. but there is a problem which reducing the cell recovery rate in this method and in most cases, it causes unsuccessful isolation. We see the tissue fragments after culture, frequently float up and lose their connection from the bottom of the culture dishes.

To improve extraction of MSCs from the umbilical cord by the explant method, we used sterilized metallic meshes to fix the chopped tissues on the bottom of the plates.

The plates were pre-coated with 0.1 mg/mL Poly-L-lysine and 1% gelatin. After 3-5 days of culturing and migration of cells from the pieces of tissue, a fresh culture medium was added with a ratio of 50:50 to the old culture medium in the plates (see Fig. S2).

##### 2.2.1.3. Immunophenotyping

To confirm MSC identity, they were immunophenotyped using anti-CD14, CD45, CD34, CD44, CD90, and CD105 antibodies conjugated with Fluorescein isothiocyanate (FITC) and Phycoerythrin (PE). The cells were trypsinized after the third passage, incubated with an individual antibody for 45 min at 4 °C, and then analyzed using a flow cytometer. The data was then analyzed using FlowJo software.

##### 2.2.1.4. Colony forming unit (CFU) assay

To measure MSCs’ spontaneous reproduction capacity in the first passage, the cells were cultured at a density of 2 cells per cm^2^ in a plate and incubated for 10-14 days. The cell colonies were then stained using 2% crystal violet, and imaging was done with phase contrast microscopy.

##### 2.2.1.5. Optimization of conditions for expanding MSCs and collecting the Secretome

The cells were sub-cultured between passages 3 and 7 using T125/T175 cell culture flasks. Once they reached 80-90% confluence, the culture media was replaced with FBS-free media. Following a 48-hour incubation, the cell media was collected and centrifuged at 2000 g for 20 minutes at 4 °C. The supernatant was then transferred to a new tube and subjected to a second centrifugation at 10000 g for 30 minutes at 4 °C. Next. The supernatant was filtered through a 0.2 µm membrane to isolate the Secretome. The resulting Secretome was aliquoted, flash-frozen using liquid nitrogen, and stored at -70°C until further tests.

The secretome collection method was modified to enhance MSC survival and obtain a potent extract .Over three days, the serum concentration in the medium was gradually reduced from 20% to 1%. And then it was completely removed. Additionally, DMEM High glucose was replaced with RPMI-1640 media in a ratio of 80% RPMI to 20% DMEM. Subsequent steps for secretome collection followed a similar procedure as described above.

#### 2.2.2. αSN-AGs Production

##### 2.2.2.1. αSN expression, purification, and fibrillation

Human αSN cDNA was transformed into Escherichia coli Strain BL21 (DE3) (Novagen, Madison, Wis., USA) using the pNIC28-Bsa4 plasmid. Protein production was induced at OD_600_ 0.6-0.8 with 0.5 mM IPTG. After 16 h, αSN was extracted and purified osmotic shock, anion-exchange chromatography, and size exclusion chromatography (Morshedi *et al*., 2015). Absorption at 280 nm was measured to determine protein concentration with a NanoDropTM 1000 spectrophotometer (Thermo Scientific) using a theoretical extinction coefficient of 0.412 mg^−1^ mL^−1^ cm^−1^. Purified αSN was dissolved at 70 µM in a fibrillation buffer (1X PBS with 0.2 mM PMSF, 1 mM EDTA, and 0.05 mM NaN_3_). After centrifugation at 2000 g for 10 minutes, αSN was incubated in a 96-well plate with 3 mm diameter glass beads for 7 and 24 hours at 37 °C with 300 rpm orbital shaking.

##### 2.2.2.2. ThT fluorescence intensity measurement

Thioflavin T (ThT) binds to the β-sheet structures found in amyloid fibrils with an accompanying increase in fluorescence. To monitor fibril formation, 20 µL of 7 h and 24 h-aged αSN-AG was mixed with 980 µL of ThT solution (30 μM in 10Mm Tris buffer, pH 8.0). The fluorescence intensity was then measured using a Varian Cary Eclipse fluorescence spectrophotometer (Mulgrave, Australia) with excitation at 440 nm, while emission was recorded from 450 nm to 600 nm with slits set at 5 and 10 nm, respectively.

#### 2.2.3. Cell cultures

##### 2.2.3.1. hCMEC/D3 culture

Human Cerebral Microvascular Endothelial cell line (hCMEC/D3) is a well-known model for BBB studies (Poller et al.,2008). The cells were cultured in RPMI-1640 media supplemented with 5% FBS and incubated at 37 °C in a humidified incubator (with 5% CO_2_ and 90% humidity).

##### 2.2.3.2. Assays on the impact of αSN-AGs on hCMEC/D3 cells in the absence or presence of MSCs-secretome

Several assays were conducted to investigate the potentially toxic effects of αSN-AGs (7 h and 24 h-aged) on hCMEC/D3 cells, including MTT, NO production, apoptosis, and fluorescent staining. Additionally, we examined the impact of secretome-derived UC-MSCs on hCMEC/D3 cells when exposed to αSN-AGs. To perform these experiments, hCMEC/D3 cells were cultured at a density of 10^4^ cells per well in 96-well plates coated with 0.1% type I collagen. Upon reaching 90% confluency, the cells were treated with different αSN-AGs (10% v/v, 7 h and 24 h-aged) either in the absence or presence of the Secretome at 100 and 400 µg/mL for 24 hours.

###### 2.2.3.2.1 Cell Metabolic Activity Assessment

MTT assay explored the impact of Secretome on hCMEC/D3 cell survival and αSN-AGs-induced metabolic toxicity at different concentrations. HCMEC/D3 cells were cultured at 2×10^4^ cells/well in 96-well plates at 90% confluence. Various concentrations of the Secretome (ranging from 10μg/mL to 1000μg/mL) were administered to the culture. To study the impact of Secretome on αSN-AG toxicity, cells were exposed to 7, 24, and 48h-aged αSN-AGs. Secretome was added at 100 µg/mL concentrations and 400 µg/mL. The cells were then incubated for 24 hours. Each well was supplemented with 50 µg/mL MTT in PBS. The cultures were subsequently incubated in the dark at 37 °C for 4 hours, after which the formazan crystals formed were dissolved using DMSO and the absorbance of the resulting solution at 570 nm measured on a microplate spectrophotometer (Epoch 2, BioTek company, Gen5 software, USA).

###### 2.2.3.2.2 Nitric oxide (NO) production assay

Reactive nitrogen species (RNS) are generated due to regular cell functioning, similar to reactive oxygen species (ROS). Apart from their vital functions as signaling molecules in biological processes, RNS have the potential to induce oxidative stress. The predominant RNS is nitric oxide (NO), produced when L-arginine is oxidized to L-citrulline through NADPH-dependent NO synthetase. First, hCMEC/D3 cells were treated with Secretome (100-2500 μg/mL) to investigate the secretome effect on hCMEC/D3 Cells. After that, these cells are treated with αSN-AGs in the absence or presence of a secretome. The production of NO from hCMEC/D3 cells was explored using the Griess test. The cells were cultured in a 96-well plate and, after 24 hours, treated with αSN-AGs and Secretome (100 µg/mL and 400 µg/mL) and then incubated for more than 12-15 hours. Following this, 50 μL of cell supernatant was combined with 50 μL of 1% sulfanilamide and incubated for 10 minutes. Afterwards, 50 μL of 0.1% N-(1-naphthyl) ethylenediamine (NED) was added, and the absorbance at 540 nm was recorded. A NO standard curve was generated by creating a series of dilutions from a nitrite solution ranging from 0 to 200 μM of nitrite to determine the quantity of NO generated in each well.

###### 2.2.3.2.3. Morphological Analysis of Cells

The cells were grown in a 96-well plate pre-coated with type I collagen at a density of 15,000 cells per well. After 24 hours of incubation, the cells were exposed to 7-h and 24-h aged αSN-AGs (10% volume per volume) and 100 μg/mL of Secretome. They were then incubated for another 24 hours. The cells were rinsed three times, fixed with 3.7% paraformaldehyde, permeabilized with 0.2% Triton X-100, and stained with DAPI and ThT. In each well, a concentration of 250 μM of ThT was added. After 30 minutes of incubation in darkness, the ThT solution was removed, and the cells were washed with PBS. Next, DAPI was added to the wells at a one µg/mL concentration and incubated for 5 minutes at room temperature. Subsequently, the cells were washed three times with PBS and then mounted for imaging. Finally, imaging was performed using a Ceti inverso TC100 fluorescence microscope.

###### 2.2.3.2.4. Flow cytometry assay procedure

We examined the cell apoptosis and necrosis level in hCMEC/D3 cells when exposed to αSN-AGs and Secretome. The cells were grown in a 6-well plate coated with type I collagen at a density of 3×10^5^ cells/well and incubated for 24 hours. Subsequently, the cells were treated with a mixture of 7-hour and 24-hour-aged αSN-AGs (10% (v/v)) and Secretome (200 μg/mL) and incubated for 48 hours. After this, the cells were detached using trypsin and suspended in 500 µL of binding buffer. FITC labeled with Annexin V and Propidium Iodide (PI) was used to stain the cells, which were then incubated in the dark for 15 minutes. The rate of early apoptosis and late apoptosis/necrosis was quantified using a BD FACS Calibur flow cytometer, and the data obtained were analyzed using FlowJo software v.7.6.1.

###### 2.2.3.2.5. Scratch wound assay to assess wound healing activities of hCMEC/D3 cells

The cells were grown in a 24-well plate with a density of 5×10^4^ and incubated for 24 hours. Subsequently, a straight scratch was created using a 100 µL sterile tip. The cells were then rinsed twice with PBS and treated with serum-free media supplemented with 5% (v/v) αSN-AGs and various concentrations of Secretome (50, 100, and 400 μg/mL). They were incubated for 48 hours. Afterward, the migration of cells into the empty region was assessed by measuring the filled area using Digimizer and ImageJ software, employing the following equation (1).

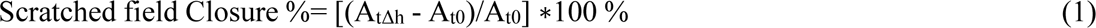

A_t0_ = the area of the scratch wound at time zero

A_tΔh_= The area of the scratch wound at time h

##### 2.2.3.2. In vitro model of the BBB

###### 2.2.3.2.1. Monoculture model

hCMEC/D3 cells were cultured on a transwell-24 well plate with a 0.4 μm pore polyester membrane insert. The insert was pre-coated with 0.1% collagen type I. The cells were seeded at a concentration 3×10^4^ cells/cm^2^ in RPMI-1640 culture media containing 5% FBS, 2 mM GlutaMAX, and 1.4 μM hydrocortisone. The culture was incubated at 37 °C, 5% CO_2_, and 90% humidity. The cells’ TEER (transendothelial electrical resistance) was measured every day starting from 24 hours after cultivation. When the TEER reached a stable value, it indicated the formation of a monolayer model, which can be considered a working model of the BBB (see Fig S9).

###### 2.2.3.2.2. Co-culture of MSCs with hCMEC/D3

UC-MSCs were cultured at a density of 6×10^3^ cells/cm^2^ in the bottom of a transwell-24 well plate that had been pre-coated with 0.1% collagen type I. They were incubated for 24 hours. Following this, hCMEC/D3 cells were cultured on the luminal side of a transwell insert with a pore size of 0.4 μm and a polyester membrane. The density of hCMEC/D3 cells was 3×10^4^ cells/cm^2^ The hCMEC/D3 cells were co-cultured with the UC-MSCs in indirect contact until the final day of treatment.

###### 2.2.3.2.3. Transepithelial/endothelial electrical resistance (TEER) assay

To assess the integrity of our BBB models, the TEER value (Ω/cm) was continuously measured using EVOME2 (World Precision Instruments, USA). After 5 to 7 days, when the TEER value stabilized, the cells were exposed to different αSN-AGs (7 h and 24 h-aged, 10% (v/v)), with or without Secretome (35% (v/v)). Subsequently, the TEER value was recorded every 24 hours for 4 days by replacing the upper culture medium of the insert with a fresh medium. The final TEER was determined by subtracting the TEER of each well from the TEER of the blank (empty insert containing culture medium) and multiplying it by the area of the insert (0.33 cm^2^) using Eq. (2). In the co-culture model, the TEER of hCMEC/D3 cells was also measured using the same method, with the exception that the inserts containing hCMEC/D3 cells co-cultured with UC-MSC were placed in wells containing culture medium without UC-MSCs.

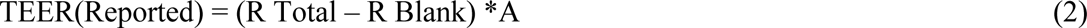

R Total = Total resistance (Ω) of the cell layer

R Blank = The resistance of the coated blank well (without cell)

A = The surface area of the filter (0.33 cm^2^)

###### 2.2.3.2.4. BBB permeability measurements

The hCMEC/D3 cells cultured as a monolayer on transwell inserts were exposed to 10% (v/v) of 7-h and 24-h aged αSN-AGs with or without Secretome (35% (v/v)) for 48 hours. After incubation, the media in the upper chamber of the transwells were replaced with fresh media containing 1% FBS and FITC-Dextran at a final concentration of 100 μg/mL. The media in the lower chamber of the wells were replaced with 600μL of serum-free fresh media. Subsequently, the media in the lower insert were collected and replaced with fresh media at 30-minute intervals and at 1, 3, 5, 7, 12, and 24 hours. The FITC fluorescence intensity was measured at an excitation wavelength of 495 nm (slit 5) and an emission wavelength of 525 nm (slit 10). The permeability (Pe) of FITC-Dextran was determined using the apparent permeability (Papp) and a standard curve following the method mentioned earlier (Hourfar *et al*., 2023).

###### 2.2.3.2.5. Analysis of the Release of Cytokines and Chemokines

The UC-MSCs were grown in the bottom of 24-well plates at a density of 6×10^3^ cells/cm^2^. After 24 hours, hCMEC/D3 cells were grown on the luminal side of the transwell-24 well plate pre-coated with 0.1% collagen type I at a density of 3×10^4^cells/cm^2^. Once the hCMEC/D3 cells had formed a complete monolayer on the transwell insert, they were treated with αSN-AGs (10% v/v). To serve as a control, hCMEC/D3 cells were grown without UC-MSCs. After 12, 24, and 48 hours of treatment, samples of the upper and lower parts of the insert were collected (100 µL each) to measure their cytokines and chemokines content. ELISA assays were used to analyze IL-10, IL-8, TNF-α, and human CCL2 levels, representing both inflammatory and non-inflammatory chemokines and cytokines. The absorbance of samples and standard at 540 nm was recorded using Hyperion MPR4++ Microplate Reader.

## Statistical analysis

The data were collected in triplicates and presented as the mean ± standard deviation (SD). Statistical analysis was performed using SPSS software version 25.0. One-way ANOVA was used to determine the presence of statistically significant differences between groups. A significance level of **p ≤ 0.01 and *p ≤ 0.05 was considered significant.

## 3. Results

### 3.1. Extraction of Highly Purified Mesenchymal Stem Cells (MSCs) from the Wharton’s jelly and and Placenta junction of Umbilical Cord

*In vitro*, MSCs are characterized by spindle-shaped morphology, adhesion to the surface, self-renewal, formation of cell colonies, and the expression of specific markers on their surface (Haasters *et al*., 2009). After extraction from the human umbilical cord using the modified explant method (Fig. S1, S2, S3). A modified-explant method was used to increase the efficiency of isolating MSCs from umbilical cord. The pieces of umbilical cord tissue until the 14^th^ day (before the first passages) were attached to the bottom of the plate by sterilized metallic meshes, which helped the mesenchymal stem cells to migrate out of the tissue with high potential.

MSCs were analyzed after the third passage. The results showed that these cells have the expected spindle-shaped morphology (Fig.1a) and have the ability to self-renewal and form cell colonies (Fig.1c). They also expressed specific markers of human MSCs (CD44, CD90, CD105) on their surface. Gratifyingly, they were negative for non-specific markers belonging to endothelial cells (CD34), monocytes and macrophages (CD14), and hematopoietic cells (CD45) (Fig. 1b). Therefore, the extraction of MSCs from human umbilical cord tissue had a purity of over 99% and was free of cross-contamination.

**Figure 1.**
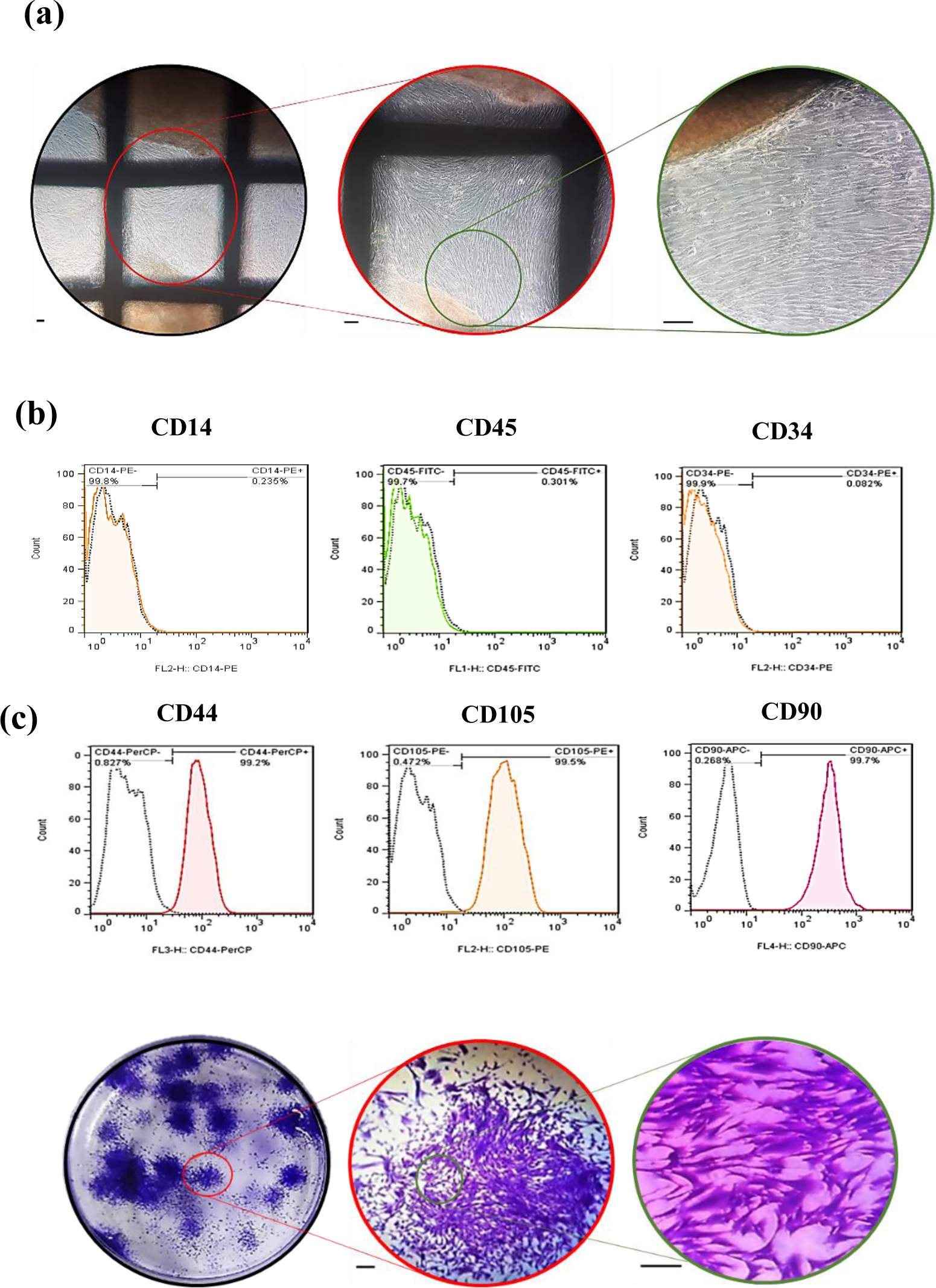
Characterization of MSCs derived from umbilical cord using a modified-explant method. (a) UC-MSCs before the first passages had fibroblast-like morphology. Scale bar = 200 μm (b) Flow cytometry analysis showed stained hUC-MSCs in third passages were positive for mesenchymal markers (CD44, CD105, and CD90) but not for non-mesenchymal markers (CD34, CD14, CD45). (c) Colony-forming capacity. UC-MSCs were cultured at a density of 2 cells/cm^2^ in Petri dishes in first passages. After 21 days, single colonies of cells were stained with 0.1% crystal violet. Scale bar = 100 μm

### 3.2. Selecting and extraction from UC-MSCs condition media

Changes in the condition of MSC culture media directly impact the Secretome’s characteristics and contents. Under serum-deprived conditions, MSCs modify their secretory content, enhancing the abundance of secretory factors and extracellular vesicles. These alterations hold the potential for therapeutic applications. Previous studies have proven that the culture media for MSCs, which lacks serum and nutrients, enhanced the release of therapeutic factors within the conditioned media (Haraszti *et al*., 2019). In this study, we used two different serum-free culture media: RPMI and RPMI supplemented with DMEM-High glucose. The results indicate that hUC-MSCs displayed high viability and the lowest nitrite production levels after being cultured in a mixture of RPMI and DMEM-High glucose (80:20 (v/v)) media for 24 and 48 hours (Fig. 2a, b).

**Figure 2.**
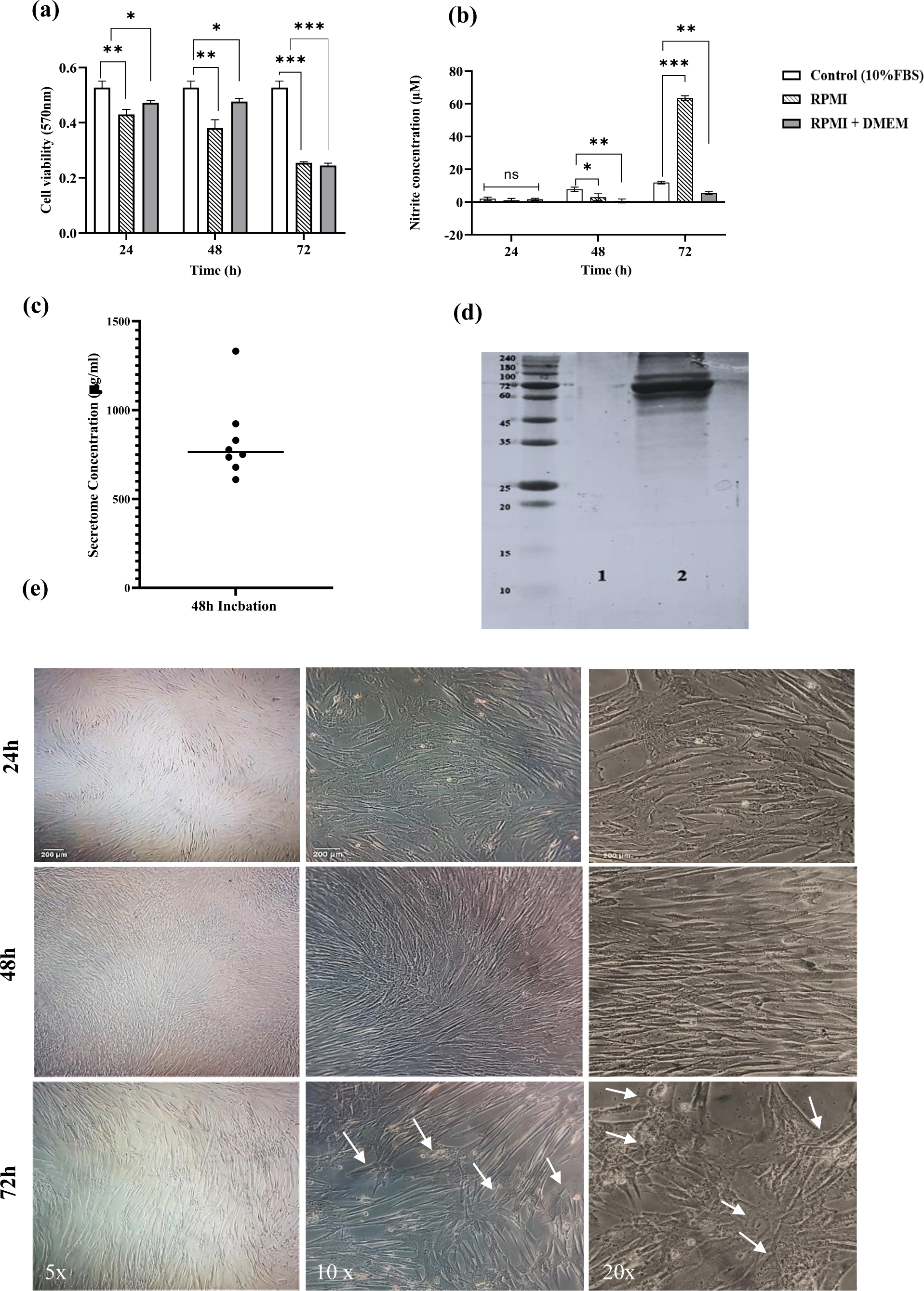
Selecting and analysis of the proper conditions for collecting hUC-MSCs CM. (a-b) The effect of the different cell culture mediums on hUC-MSCs viability and nitric oxide release was measured. UC-MSCs were cultured in serum-free media (RPMI-1640 or DMEM-Hg: RPMI with an 80:20 ratio) after 24-hour pre-culturing. (c) the concentration of protein content of the isolated secretome was measured using BCA kit. (d) The protein allocation was assessed using SDS-PAGE, line 1: consisting of RPMI supplemented with DMEM-High glucose (80:20 (v/v)), line 2: the CM collected from the UC-MSCs after 48 hours. (e) The morphology of UC-MSCs was observed using inverted microscopy after 24, 48, and 72 hours culturing with three magnifications (5×,10×,20×). The white arrows show the damaged and apoptotic cells that appeared after 72 hours of incubation to collect the CM. but no visible effects on cell morphology were observed after 48 hours of incubation. Statistical analysis was performed using one-way ANOVA with a Tukey post-hoc test (*P-value ≤0.05, **P-value ≤0.01, ***P-value ≤0.001, N = 3, Mean ± SD).

In addition, we analyzed the Secretome isolated after 48 hours to assess its content and protein concentration. This analysis was performed using a BCA protein concentration assay and SDS-PAGE (Fig. 2c, d), which shows a high protein content in the secretome band (average value of 720 μg/mL) compared to culture media. Additionally, we examined the morphology of UC-MSCs at 24, 48, and 72 hours’ post-collection of the conditioned medium of RPMI-DMEM. The cells only showed insignificant changes in their morphology after 48 hours of media collection; after 72 hours were more clearly visible (Fig. 2 e).

### 3.3. UC-MSCs’ Secretome protects brain capillary endothelial cells against αSN cytotoxicity

The initial investigation assessed the impact of UC-MSCs Secretome on metabolic activity in the hCMEC/D3 cell line using MTT and Griess assay (Fig. 3a-b). Subsequently, hCMEC/D3 cells were co-treated with Secretome and αSN-AGs (Fig. 3d-i). Our findings revealed that the Secretome, at concentrations up to 1 mg/mL, had no significant influence on cell viability (Fig. 3a). Concentrations up to 2.5 mg/mL did not affect the production of nitrite species in hCMEC/D3 cells (Fig. 3b). Both 7h and 24h αSN-AGs caused a significant decrease in cell viability and an increase in the production of toxic nitrite species in hCMEC/D3 cells (Fig. 3d-i). However, treatment with UC-MSC secretome at concentrations of 100 µg/mL (Sec I) and 400 µg/mL (Sec II) effectively prevented toxicity induced by 7h and 24h αSN-AGs (Fig. 3d, e). However, the Secretome did not protect against 48h αSN-AGs (Fig. 3f). Additionally, the Secretome reduced the levels of nitric oxide (NO) caused by αSN-AGs in the supernatant of hCMEC/D3 cells (Fig. 3h, i). Remarkably, > 60% of the cells underwent apoptosis in response to αSN-AGs, but this percentage significantly decreased after treatment with Secretome (Fig. 3j-l and Fig. S6).

**Figure 3.**
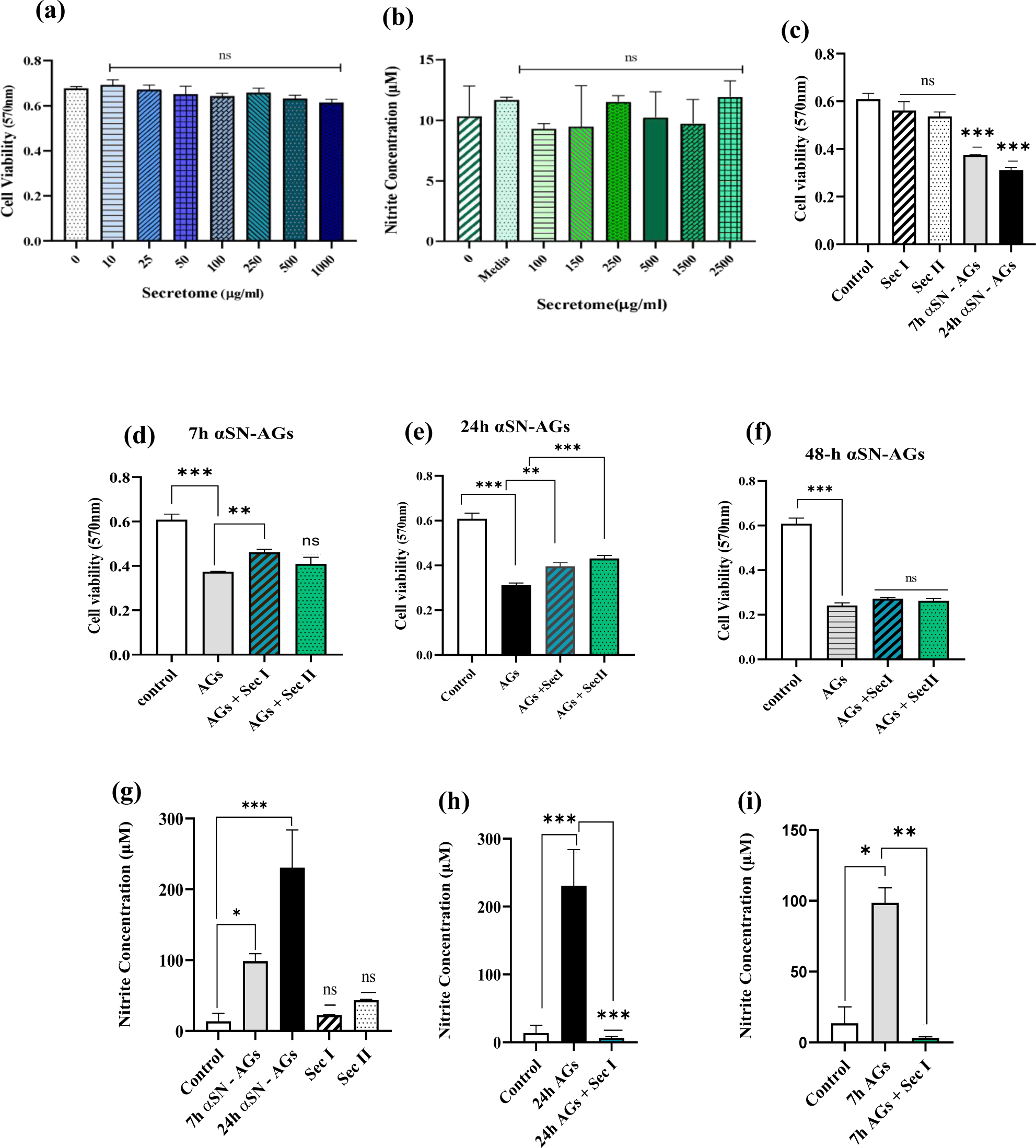
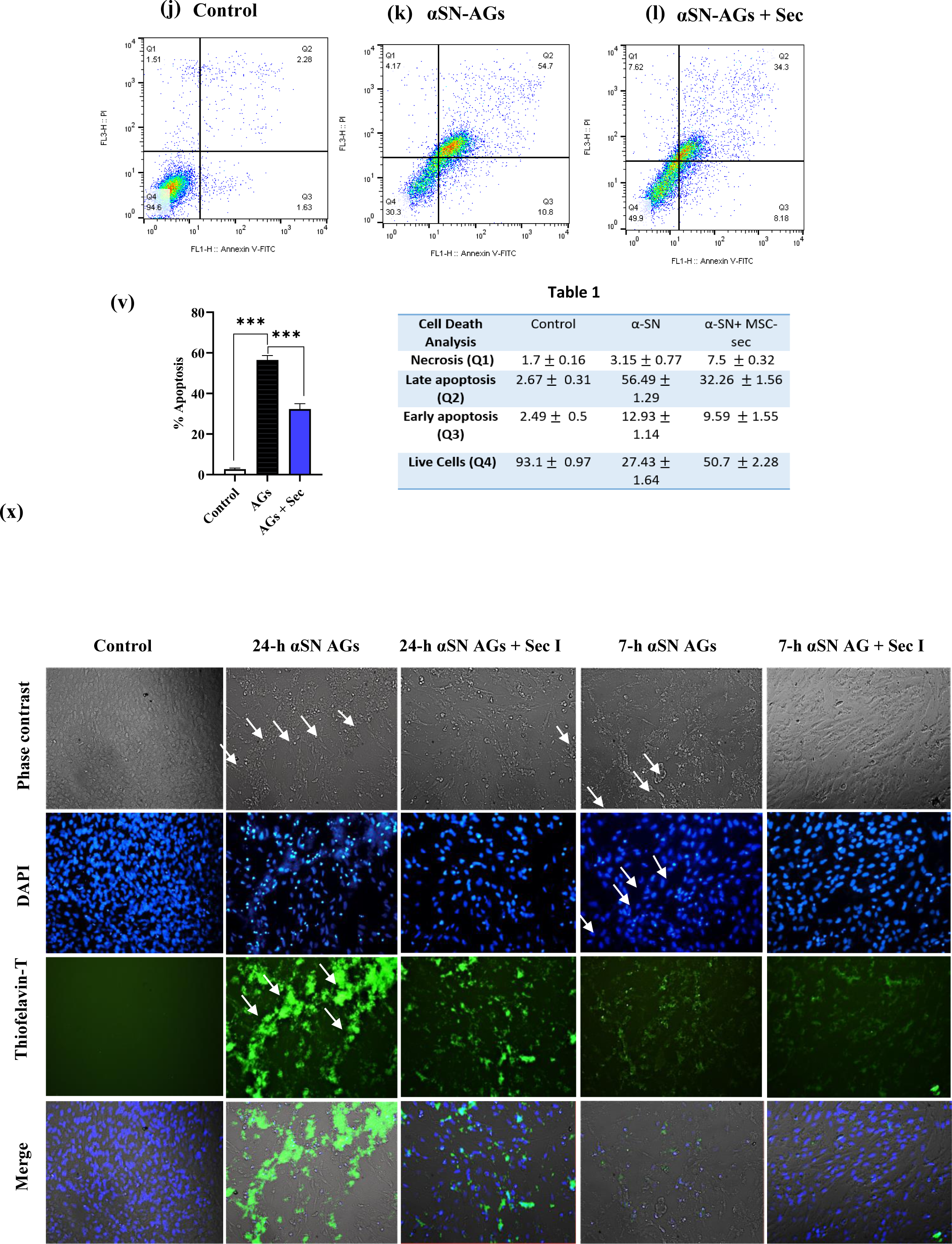
The protective effects of hUC-MSCs secretome on hCMEC/D3 cells against αSN-AGs. The impacts of the secretome were evaluated by assessing cell viability, NO production, apoptosis, and immunofluorescence staining. (a-b) Cell viability and nitrite level in hCMEC/D3 cells after 24 hours of treatment with different concentrations of secretome derived from hUC-MSCs (c)The viability of hCMEC/D3 cells in the presence of different species of αSN-AGs; (d) 7h αSN-AGs, (e) 24h αSN-AGs, (f) 48h αSN-AGs, with or without hUC-MSCs secretome at two different concentrations of 100 μg/ml (Sec I) and 400 μg/ml (Sec II). (g-i) the levels of nitrite in the supernatant were measured after treatment of (g)7h αSN-AGs (h) 24h αSN-AGs and (i) 48h αSN-AGs on the hCMEC/D3 cell culture in the presence and the absence of hUC-MSC secretome. (j-v) Exploring the effect of αSN-AGs on the cell apoptosis in the presence or the absence of hUC-MSCs secretome. The representative histogram shows viable cells (Q4, double negative), early apoptotic cells (Q3, Annexin V-FITC positive), late apoptotic/necrotic cells (Q2, Annexin V-FITC, and PI positive), and necrotic cells (Q1, PI-positive) in (j) control cells, (k) 24h, 48h, 7h -aged αSN-AGs in equal ratio treated cells (15% w/w) (l) αSN-AGs co-treatment with hUC-MSCs secretome (200 μg/ml) on the hCMEC/D3 cell lines. Table 1 provides the percentage of the cells in each quadrant as mean ± SD. (v) Quantitative measurement of apoptosis percentage in hCMEC/D3 cell line after 48 h incubation in the presence of αSN-AGs with or without hUC-MSC secretome. (x) Immunofluorescence staining analysis of hCMEC/D3 cells after treatment with 7h, 24h αSN-AGs with or without the hUC-MSCs-derived secretome. hCMEC/D3 illustrated with DAPI-stained nuclei (blue color) and Thioflavin-T (green color) for staining αSN-AGs. Nucleus fragmentation leading to apoptosis was observed in 24h and 7h αSN-AGs treatment groups (White arrows). Also, the intensity of ThT-fluorescence in this group increased (White arrows). In contrast, in treatment with hUC-MSCs secretome at 100 μg/ml, the intensity of AGs and nucleus fragmentation decreased. *P-value ≤0.05, **P-value ≤0.01, ***P-value ≤0.001 (N = 3, Mean ± SD), one-way ANOVA, Tukey post-hoc test.

The quantitative data on apoptosis is presented in a graph (Fig. 3v) and Table 1. Additionally, fluorescent images confirm the earlier findings, revealing fragmented and apoptotic nuclei through DAPI staining. ThT staining confirms the aggregation intensity and cellular damage observed after treatment with 24h αSN-AGs. Interestingly, the presence of Secretome in both the 7h and 24h aggregated species groups mitigates the severity of the injury (Fig. 3x).

### 3.4. hUC-MSCs Secretome Aids Healing and Alleviates Dysfunction in Brain Capillary Endothelial Cells due to αSN-AGs

A scratch wound assay also known as the scratch assay was employed to investigate the impact of the hUC-MSC Secretome on the migratory capacity of hCMEC/D3 cells. αSN-AGs influence the migration of hCMEC/D3 cells by impeding regular migration or causing aberrant individual cell migration (Hourfar *et al*., 2023). Following scratching, the cells underwent treatment with hUC-MSCs-derived Secretome at varying concentrations (50, 100, and 400 µg/mL) in a media devoid of serum (Fig. 4a). The findings demonstrated that the Secretome prompted cell migration in all concentrations, particularly at 100 μg/mL Subsequently, to examine the impact of Secretome derived from hUC-MSCs on migration in the presence of AGs, the hCMEC/D3 cells were simultaneously exposed to a concentration of 100 μg/mL of Secretome and 24h αSN-AGs. As depicted in (Fig. 4b) where αSN-AGs were used to treat the sample, the hCMEC/D3 cells migrated individually and sporadically but suffered significant cell death. However, when αSN-AGs were combined with Secretome, the areas without cells in the scratch site and behind it was reduced. This suggests that the Secretome prevented cell death during migration, promoted cell migration, and repaired the scratch site in the presence of αSN-AGs compared to the control group.

**Figure 4.**
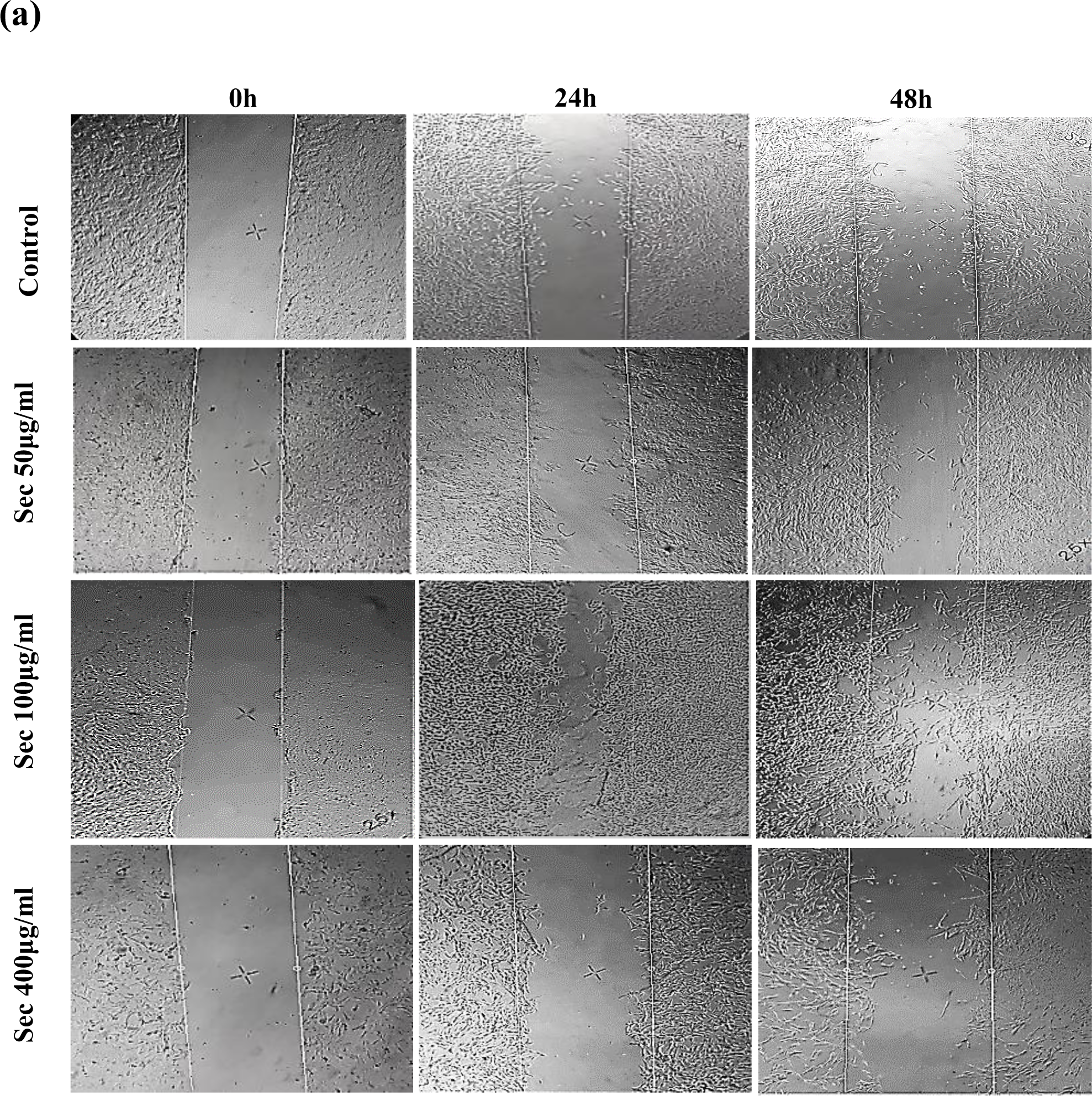
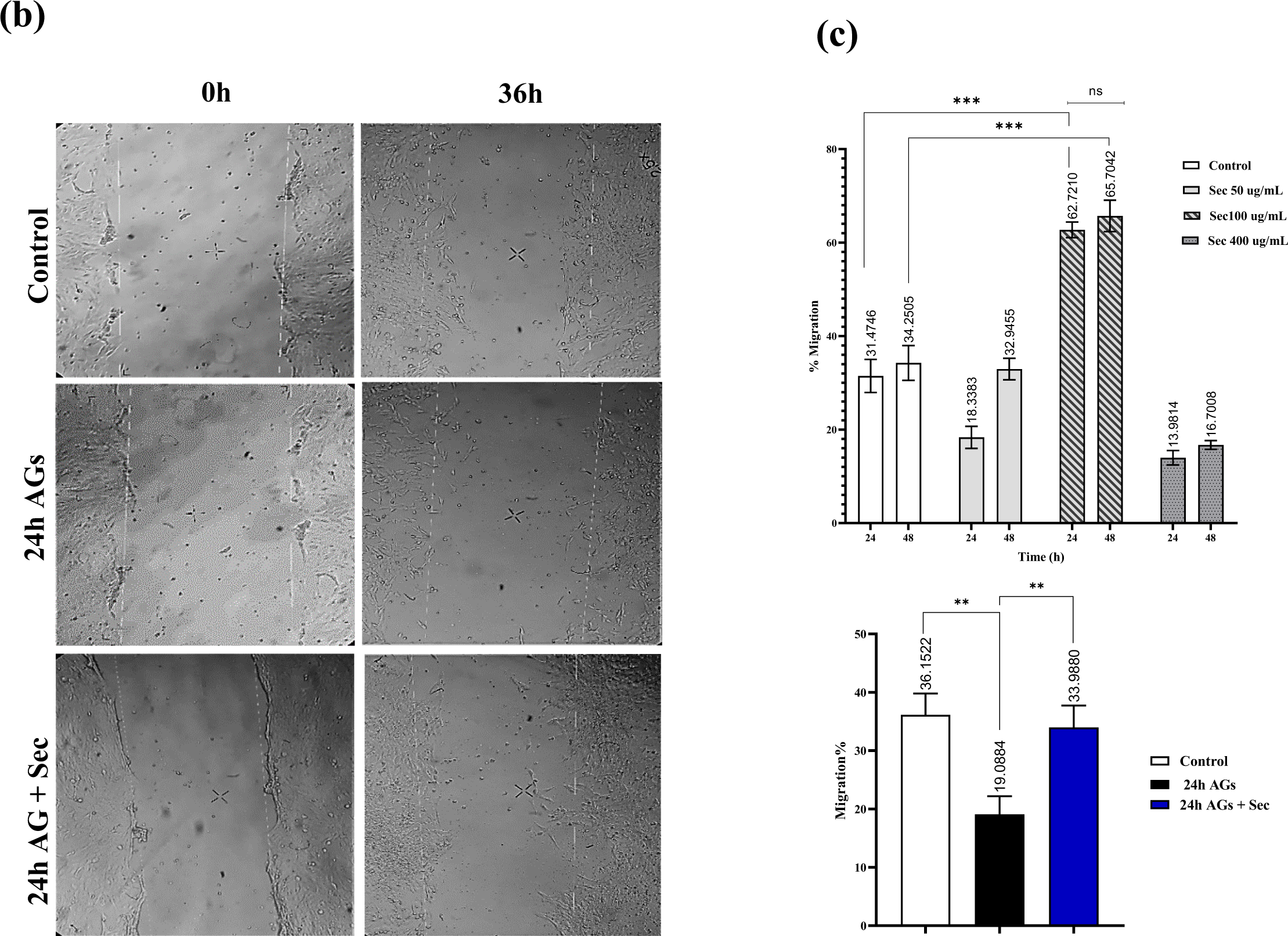
Use of *in vitro* wound healing assay, to assess the effect of different doses of UC-MSC Secretome on wound healing, with or without αSN-AGs. (a) The hCMEC/D3 cells were cultured until they reached 90% confluence, then a scratch was made and the cells were treated with MSC-secretome at various concentrations (50, 100, 400 μg/ml). The closure of the scratch area within 48 hours was observed using phase contrast microscopy and analyzed using ImageJ and DigiMaizer software. (b) The wound healing assessment was performed on hCMEC/D3 cells treated with 24h αSN-AGs (5% w/w), with or without secretome (100 μg/ml), in FBS-Free Media for 36 hours. (c) and (d) The quantitative results of wound healing extracted from part (a) and (b), respectively. *P-value ≤0.05, **P-value ≤0.01, ***P-value ≤0.001 (N = 3, Mean ± SD), one-way ANOVA, Tukey post-hoc test

### 3.5. Enhancing BBB Integrity: The Protective Effect of hUC-MSCs-Derived Secretome on αSN-AGs Induced damage in an *In Vitro* BBB Model

To assess the impact of the Secretome derived from hUC-MSCs on the functionality of the BBB, we utilized an in vitro model of the BBB (see Figure S9). The hCMEC/D3 cells were treated with αSN-AGs or the Secretome by placing them on a transwell. Subsequently, we conducted assays to measure the TEER and permeability indexes.

The findings showed that 7h and 24h αSN-AGs decreased the TEER value over a 72h period (Fig. 5b, d) and increased the permeability index (Fig. 5a, c). However, treatment with Secretome from both groups (7h and 24h αSN-AGs) effectively prevented the decline in BBB function (Fig. 5a-d). To investigate the impact of fresh Secretome, which is directly secreted by hUC-MSCs, on the BBB impairment caused by αSN-AGs, a co-culture model was employed where hUC-MSC cells were co-cultured with hCMEC/D3 cells (with the hUC-MSCs and hCMEC/D3 cells cultured at the bottom of the well and on the apical side of the insert, respectively) (see the fig. S10). Afterward, the hCMEC/D3 cells underwent treatment with αSN AGs, and subsequently, the TEER and permeability values were measured (Fig. 5e-g). The findings demonstrated that hUC-MSCs also could mitigate the harm caused by 7h and 24h αSN-AGs (Fig 5f, g.). Based on the obtained results, it is likely that the protective effects of the Secretome on the BBB are attributed to the secretion of active factors with modulatory properties

**Figure 5.**
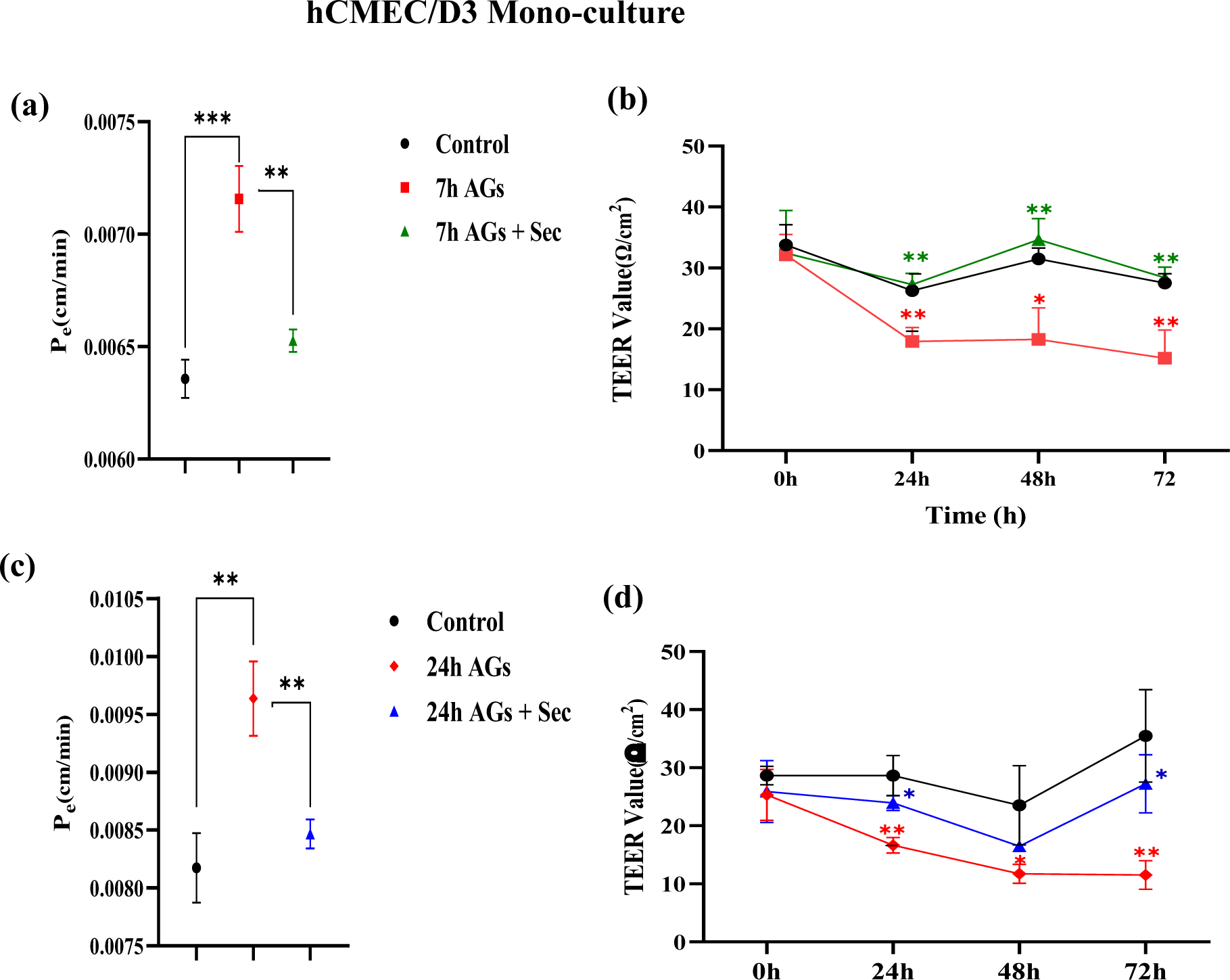
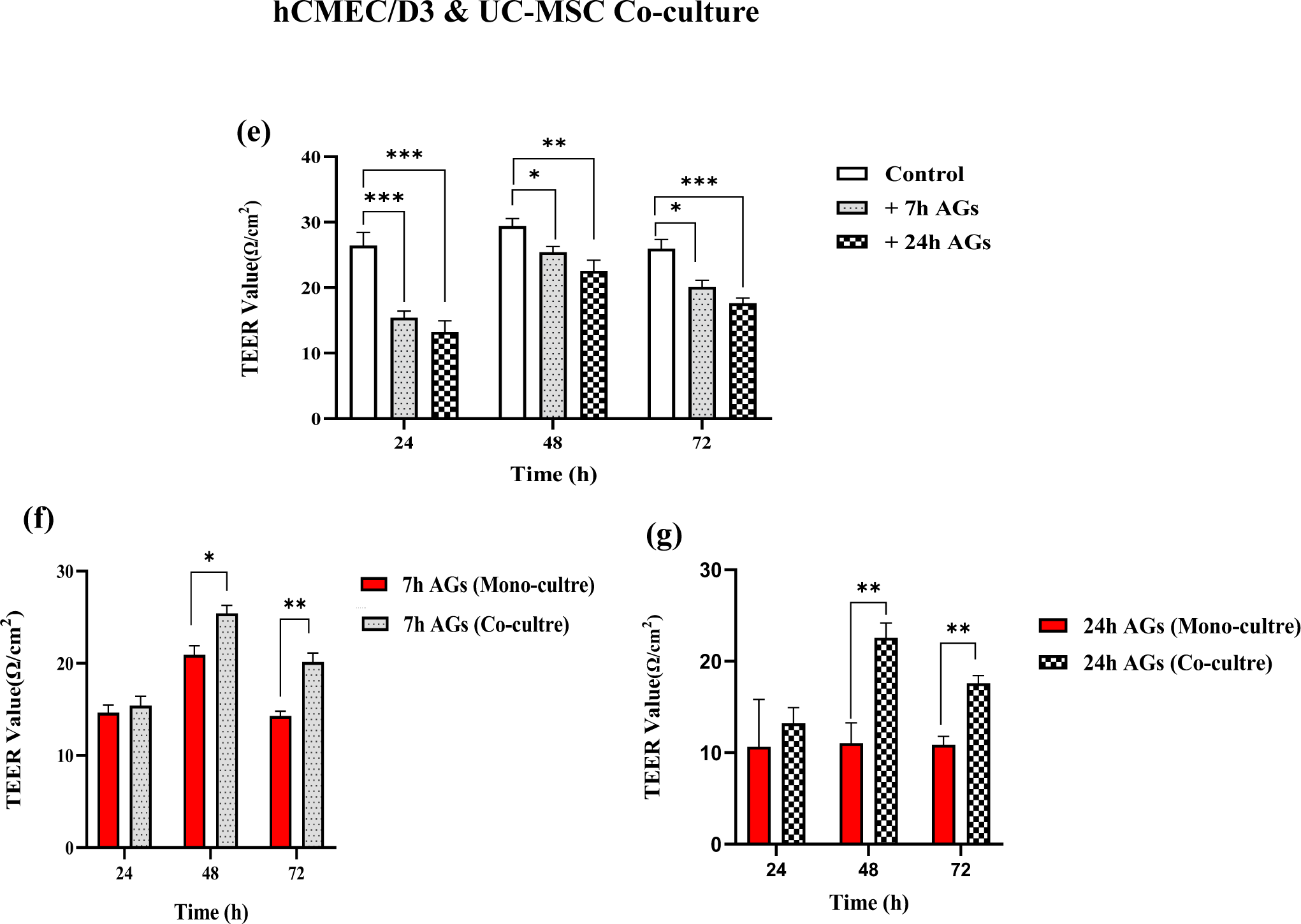
Integrity assessments of the *in vitro* BBB model in the presence and absence of 24h and 7h αSN-AGs and MSC-secretome by measurement of TEER and permeability parameters. (5a, c) Permeability assay and (5b, d) TEER measurement showed significant differences after secretome treatment in the presence of αSN-AGs in a monolayer system. (e-g) In the co-culture system, the hCMEC/D3 cell were cultured on the apical side of the transwell insert, and UC-MSC was cultured in the bottom of the 24well plate. Both cells were in communication with each other through their secretions until the end of the formation of BBB on the apical side. (e) The co-culture system was treated with 7-h and 24-h AGs (15% w/w) on the apical side, and TEER value decreased during 24, 48, and 72h. (f, g) Comparison effect of the UC-MSCs in mono and co-culture system on preventing TEER reduction after treatment with (f) 7h αSN-AGs and (g) 24h αSN-AGs. *P-value ≤0.05, **P-value ≤0.01, ***P-value ≤0.001 (N = 3, Mean ± SD), one-way ANOVA, Tukey post hoc test.

### 3.6. Potential Involvement of Cytokines and Chemokines in MSCs-Secreted Activity Against αSN-AGs Toxicity

To clarify the protective mechanism involved in MSCs-secretion on the BBB, we measured the levels of inflammatory cytokine (TNF-α) and anti-inflammatory cytokines (IL8, IL10). Our findings showed a significant increase in the level of TNF-α after 24 hours of treatment with αSN-AGs in hCMEC/D3 cells compared to the control group. However, when hUC-MSCs were co-cultured with hCMEC/D3 cells, the rise in this inflammatory factor was prevented (Fig. 6a). The findings showed that the presence of hUC-MSCs in co-culture samples resulted in a more robust expression of the anti-inflammatory cytokine IL-8. This suggests that hUC-MSCs may reduce inflammation caused by αSN-AGs damaging the BBB by releasing IL-8 (Fig. 6c). The co-culture of hUC-MSCs with hCMEC/D3 cells resulted in a significant increase in the anti-inflammatory cytokine IL-10, suggesting that hUC-MSCs may secrete this factor in response to BBB damage and protect hCMEC/D3 cells through paracrine signaling (Fig. 6b). Additionally, our findings indicated that in the in vitro BBB model, hCMEC/D3 cells produced more of the inflammatory cytokine TNF-α than CCL2 in the early hours after exposure to αSN-AGs. However, this trend reversed over time, with CCL2 secretion increasing 24 hours after treatment (Fig. 6d). Interestingly, when LPS was used as a damaging factor to the BBB, the level of CCL2 secretion was significantly higher than TNF-α, suggesting different inflammatory pathways are activated in response to these substances. Furthermore, hCMEC/D3 cells exhibited a more robust response to LPS in releasing the inflammatory factor CCL2 (Fig. S8).

**Figure 6.**
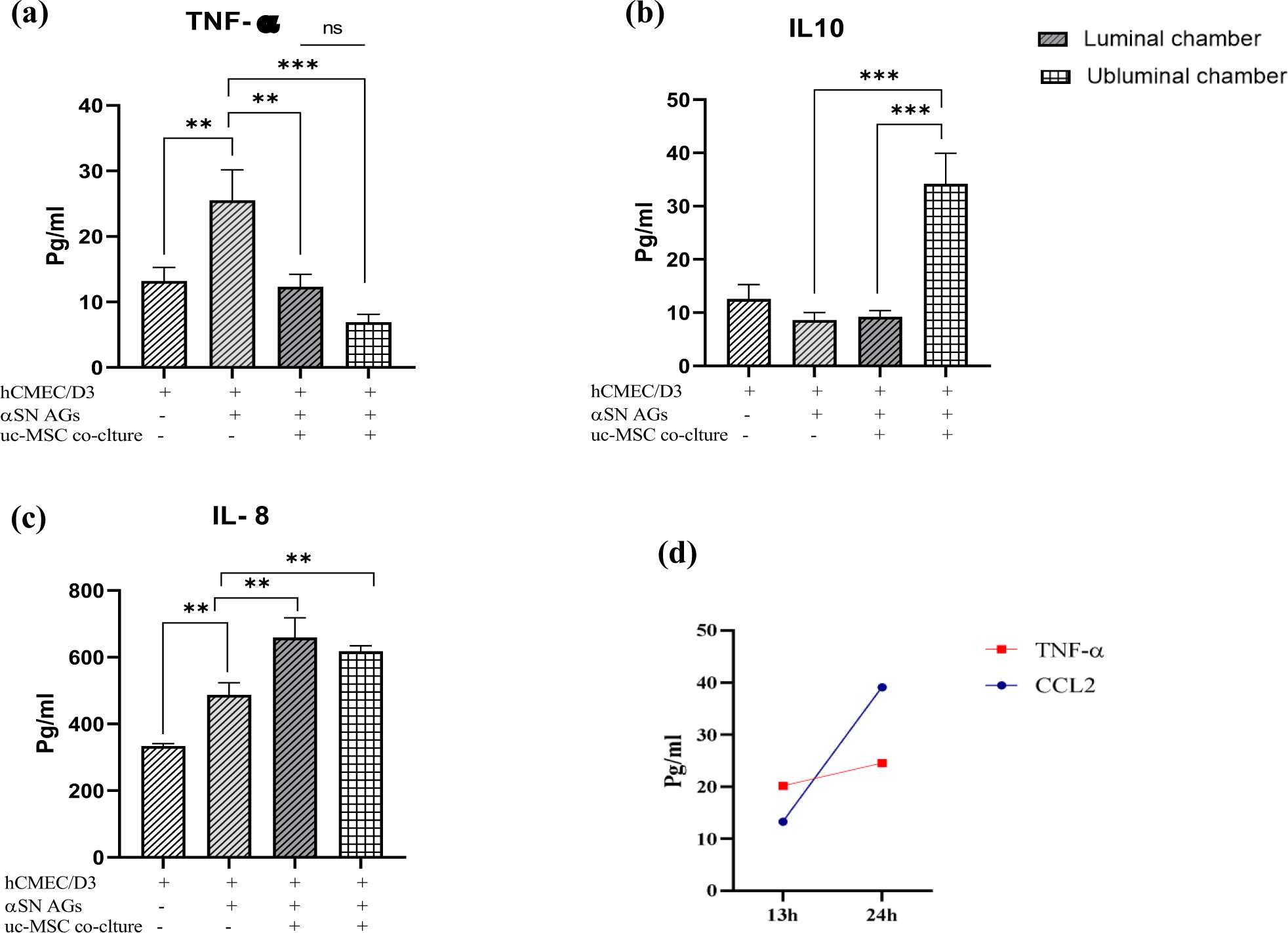
Assessment of Cytokines and Chemokines in an *in Vitro* Blood-Brain Barrier Co-Culture System. Pro-inflammatory cytokine (a)TNF-α, (b, c) anti-inflammatory cytokines IL-10, IL-8, and (e) CCL-2 chemokine were measured in co-cultured of BBB model (hCMEC/D3 culture supernatant) and conditioned media in the bottom of the well (UC-MSC culture supernatant) after treatment with αSN-AGs (15% w/w) used ELIZA assay. (d) In the monoculture model of BBB, 12 hours and 24 hours after treatment with αSN-AGs, the supernatant was collected and the amount of inflammatory factors (TNF-α, CCL2) was determined by ELISA assays.

## 4. Discussion

Previous research has indicated that the development of Parkinson’s disease (PD) is connected to damage in the blood-brain barrier (BBB) caused by activated glial cells and pericytes in response to intracellular and extracellular αSN-AGs (Konno *et al*., 2012)(Barr *et al*., 2010). This activation creates an inflammatory response in the NVU. Consequently, the BBB becomes compromised, resulting in increased permeability and the presence of inflammatory cytokines and chemokines. Furthermore, existing studies suggest that the tight junctions become permeable to blood lymphocytes, leading to BBB breakdown and the release of αSN-AGs into the peripheral blood (Schultz *et al*., 2014)(Dohgu *et al*., 2019)(Gray and Woulfe, 2015).

This study builds upon our previous research, demonstrating that the specific endothelial cells associated with the blood-brain barrier (BBB) are impacted by the Toxicity resulting from αSN-AGs. The observed effects include reduced cell viability caused by damage to mitochondria, heightened apoptosis, nitric oxide (NO) levels, and disruption of the wound healing process.

### Uc-MSC derived-secretome could protect mitochondria against αSN-AGs toxicity

Mitochondrial dysfunction is one of the most critical mechanisms of αSN-AGs toxicity in PD (Pozo Devoto and Falzone, 2017)(Gao *et al*., 2017). It has been reported that inhibition of the mitochondrial oxidative phosphorylation process increased the BBB permeability and destruction of the tight junctions (Doll *et al*., 2015). Here, by examining the mitochondrial dysfunction due to the toxicity of different αSN-AGs (7h-AGs, 24h-AGs) on the hCMEC/D3 cells, our findings revealed that the secretion of hUC-MSCs partially countered this toxicity, offering a potential preventive measure. Moreover, the apoptosis rate in the hCMEC/D3 cells treated with αSN-AGs was significantly reduced in the presence of UC-MSCs secretome. Previous studies demonstrated that MSCs, through secret trophic growth factors and cytokines, could protect the cells from mitochondria-dependent apoptosis mainly by upregulating the expression of anti-apoptotic proteins (BCL-XL, BCL-2) and downregulating the expression of apoptosis-inducing proteins (BAX, BAK, BAD) (Zhao *et al*., 2019). In addition, recent findings suggest that MSCs could transfer mi-RNAs (miR149, miR25, miR326) to the injured cell through their extracellular vesicles that regulate mitochondrial metabolism and moderate activity (Loussouarn *et al*., 2021).

### NO level decreased in hCMEC/D3 culture in presence of UC-MSC secretome

Our result demonstrated NO increase in the hCMEC/D3 culture supernatant stimulated with αSN-AGs. NO is produced in endothelial cells through the action of the nitric oxide synthase (NOS) enzyme. Endothelial cells generate NO through two forms, namely endothelial NOS (eNOS) and inducible NOS (iNOS), which contribute to various physiological functions such as regulating blood flow, ion channels, and permeability. However, during periods of stress, infections, and inflammatory conditions, NO production is induced, resulting in increased oxidative stress, apoptosis, and dysfunction of the BBB (Hurst and Clark, 1997)(Madigan and Zuckerbraun, 2013)(Thiel and Audus, 2001). given that iNOS produces NO for a longer duration (hours). In comparison, eNOS produces it for a shorter duration (minutes) (Förstermann and Sessa, 2012)(Xie *et al*., 1992); after 24 hours of αSN-AGs treatment, the increased NO levels can be attributed to the inducible type of iNOS which in turn leads to heightened apoptosis and increased BBB permeability rate.

Previous research has also established that elevated NO levels may impact the integrity and permeability of the blood-brain barrier (BBB) regarding leukocyte transmigration (Hollenberg, Guglielmi and Parrillo, 2007). In this study, the treatment of hCMEC/D3 cells with both UC-MSC secretome and αSN-AGs resulted in a notable decrease in NO. The precise mechanism by which the Secretome leads to this reduction in NO levels remains unknown. Nonetheless, substantial experimental evidence has pointed toward soluble mediators and exosomes originating from MSC as the factors accountable for this observed effect (Kemp *et al*., 2010)(Liy *et al*., 2021).

### The interaction of MSCs-derived Secretome with αSN-AGs may be involved in reducing its toxicity on BBB

Herein, we showed that MSCs-derived Secretome can prevent damage to the BBB. It has been shown that MSC cells have significant potential in protecting the mitochondria of damaged cells or even reviving this organelle in damaged cells. MSCs may accomplish this process in some ways, including establishing a balance between cleavage and fusion. By upregulating the expression of the mfn1 and mfn2 genes, MSCs induce a transition in mitochondrial shape from cleavage to fusion, enhancing respiratory parameters and safeguarding the cell (Newell *et al*., 2018).

Previous reports have also confirmed that extracellular αSN-AGs can interact with the cell membrane and attach to them. Moreover, by connecting to some receptors and extracellular channels, αSN-AGs can trigger an inflammatory response and apoptotic pathways (Surguchev, Emamzadeh and Surguchov, 2019). Hence, our findings observed that αSN-AGs can cause harm either by attaching to the cell membrane surface or infiltrating the hCMEC/D3 cells, resulting in apoptosis. Upon binding to cell surface receptors (TLR-4, TLR-2, and CD11b), αSN-AGs trigger the NF-κB apoptotic pathway (Bogale *et al*., 2021). Nevertheless, the introduction of MSCs-secretome prevented the detrimental effects inflicted by αSN-AGs. You could see an overview of our results and analysis in fig.7.

**Figure 7.**
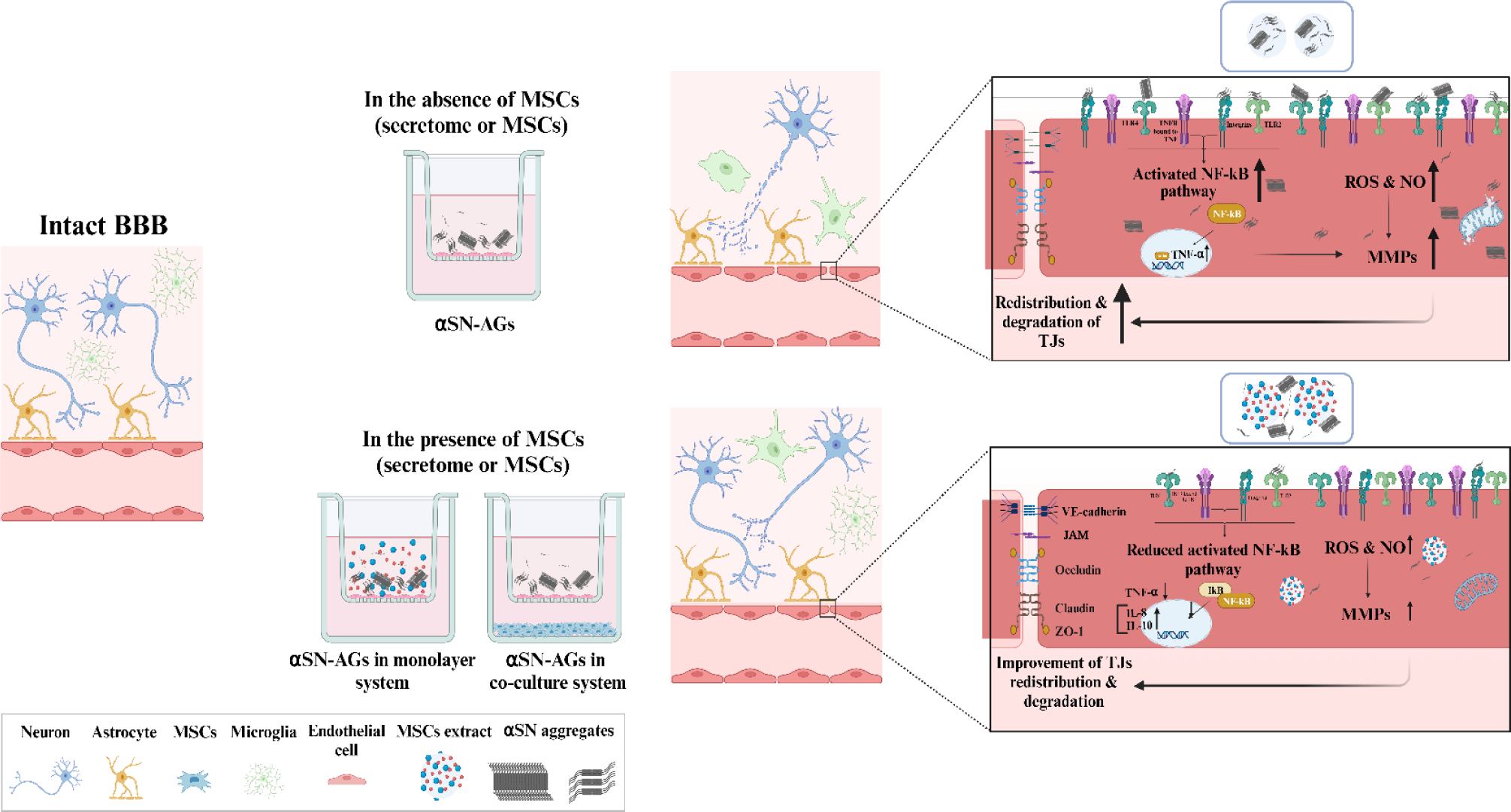
Schematic illustration of BBB during PD progression in present and absent of UC-MSCs-derived secretome and MSCs culture in an *in vitro* co-culture model. Created by https://www.biorender.com/

In addition, the contents of the Secretome (extracellular vesicles, exosomes, and proteins) may interact with different αSN-AGs species (7h-AGs and 24h-AGs), either preventing their attachment to the cell membrane or converting them to less toxic aggregates. The reduction in the ThT fluorescent intensity of αSN-AGs in the presence of Secretome may indicate this issue (Fig. S5b). We showed that hUC-MSCs-derived Secretome could interfere with the αSN-AGs through nucleation and elongation steps. Since inhibition of fibrillation is essential in the progression and prevention of the spread of PD, Secretome may prevent the progression of the disease in the early stages. Nevertheless, more investigation is needed to elucidate the mechanism behind this. There is still no detailed information about the exact mechanism of how Secretome leads to the protection of hCMEC/D3 cells or the BBB, how their contents interact with aggregated species or which parts of the secretome are most important for these protective properties.

### αSN-AGs directly induced inflammatory and anti-inflammatory cytokines and chemokines from BBB

Numerous studies have been conducted to explore the healing properties of MSCs in neurodegenerative diseases. The consensus among these studies is that the therapeutic and neuroprotective effects of MSCs can be attributed to the proteins secreted by these cells, as well as their extracellular vesicles/exosomes (Zriek, Di Battista and Alaaeddine, 2021)(Kim, Choi and Kim, 2013)(d’Angelo, Cimini and Castelli, 2020)(Rahbaran *et al*., 2022). In this regard, we aimed to investigate the potential mechanisms implicated in the protection of the BBB against damages caused by αSN-AGs. To achieve this, we examined the secretion of specific cytokines and chemokines in a BBB model treated with αSN-AGs, both in the presence and absence of hUC-MSCs secretome.

This study found that treatment with αSN-AGs caused a 2-fold increase in TNF-α secretion from hCMEC/D3 cells compared to the control group. TNF-α is an essential inflammatory factor known to be elevated in the brains of PD patients (Mogi *et al*., 1994). Previous research has demonstrated that TNF-α is the main contributor to BBB damage, and inhibiting this inflammatory factor can mitigate such damage. Interestingly, the presence of MSCs in the co-culture model reduced the extent of this damage (Zhao *et al*., 2007). This suggests that αSN-AGs may stimulate endothelial cells and increase their immunological activity. Moreover, it is possible that αSN-AGs induce the features of antigen-presenting cells (APC) in endothelial cells. The findings of this study reveal that hCMEC/D3 cells can release TNF-α when exposed to αSN-AGs. While previous research has primarily investigated the inflammatory factors produced by microglia, pericytes, and astrocytes as the primary contributors to BBB damage through inflammatory processes (Sharma *et al*., 2018)(Argaw *et al*., 2012). our results highlight the possibility of αSN-AGs playing a role in this mechanism. other investigation did not include an analysis of the direct release of inflammatory and anti-inflammatory factors from endothelial cells, except when there was a prior stimulation or pre-treatment involving at least one of the following inflammatory factors: TNF-α, IL-1β, INF-α, and IL-6 (Dohgu *et al*., 2019)(Farfara *et al*., 2019)(Argaw *et al*., 2012).

The αSN-AGs may cause apoptosis in cells by inducing the secretion of TNF-α. Our apoptotic results also may confirm increase of TNF-α secretin due to αSN-AGs treatment in BBB model.

Previous research has shown that TNF-α activates the NF-κB signaling pathway in hCMEC/D3 cells, which leads to the opening of BBB tight junctions and ultimately increases monocyte migration (Weksler, Romero and Couraud, 2013). In this regard, studies have demonstrated that beta-amyloid aggregates initiate the stimulation of this pathway within brain capillaries, consequently elevating BBB permeability (J Gonzalez-Velasquez *et al*., 2011).

### LPS induced CCL2 secretion in BBB without the mediation of blood cells

In considering investigating the ability of our *in vitro* BBB model to secrete inflammatory factors in response to external stimuli, we utilized lipopolysaccharide (LPS). LPS is known to cause loss of BBB integrity through a mechanism distinct from αSN-AGs. Our findings revealed that hCMEC/D3 cells treated with LPS showed significant secretion of the inflammatory factor CCL2 (Fig. S8), suggesting that this could be one of the mechanisms responsible for disrupting BBB integrity(Qin *et al*., 2015). According to that, the hCMEC/D3 cells in our blood-brain barrier (BBB) model secreted both TNF-α and CCL2 cytokines when treated with lipopolysaccharide (LPS) from the luminal area. Surprisingly, the amount of CCL2 secretion was nearly 10 times greater than TNF-α. CCL2, also known as MCP-1, is a crucial chemokine protein that plays a significant role in monocyte trafficking into the brain parenchyma by disrupting the BBB.

Consequently, it is involved in the inflammatory damage to the central nervous system (CNS). The elevated expression of CCL2 indicates the onset of the disease prior to the appearance of clinical symptoms (Hulkower *et al*., 1993). Similar to primary cerebral capillary endothelial cells, hCMEC/D3 cells have been observed to secrete the chemokines CCL2 and CXCL2 on both the luminal (facing the blood) and abluminal (facing the brain) sides (Subileau *et al*., 2009)(Weksler, Romero and Couraud, 2013). This supports the finding of increased CCL2 secretion in our in vitro blood-brain barrier (BBB) model after treatment with LPS. The CCL2/CCR2 signaling pathway promotes the migration of blood cells across the BBB, leading to increased permeability due to the downregulation of tight junction expression (Stamatovic *et al*., 2005). However, in our experimental design, which lacks blood cells, it appears that the decrease TEER caused by LPS damage follows a different mechanism and is not solely due to the migration of monocytes. LPS likely exerts its damaging effects through direct augmentation of permeability, while the secretion of CCL2 and recruitment of monocytes in subsequent steps contribute to the severity of the damage. Additional studies are needed to confirm this hypothesis.

### Modulation of increase in permeability and decrease in TEER in the PD-patient BBB model in the presence of MSCs secretomes may be related to the secretion of anti-inflammatory factors

In our *in vitro* blood-brain barrier (BBB) model, we found that adding αSN-AGs from the luminal side. significantly decreased trans endothelial electrical resistance (TEER) and increased permeability. One possible mechanism for the effect of αSN-AGs could be their ability to reduce the expression of tight junction proteins, particularly ZO1, and occludin, in hCMEC/D3 cells (Kuan *et al*., 2016). Additionally, a study on the intestinal epithelial barrier of Parkinson’s disease (PD) patients found a decrease in occludin expression under the influence of αSN-AGs (Clairembault *et al*., 2015). Since it is known that the expression of these proteins directly affects BBB permeability and TEER, we can assume that the observed decrease in TEER and increase in permeability induced by αSN-AGs in our study may be attributed to the reduction in tight junction protein expression.

The substantial rise of the anti-inflammatory factor IL-10 in the Secretome of MSCs suggests that these cells could safeguard hCMEC/D3 cells employing IL-10’s anti-apoptotic characteristics. Consequently, studies have demonstrated that MSC-secreted IL-10 primarily contributes to the reduction of apoptosis in pancreatic beta cells when stimulated with INF-α (Al-Azzawi *et al*., 2020). Here, we also assessed the level of IL-8. The findings indicated that when co-cultured with MSCs and hCMEC/D3 cells and treated with αSN-AGs, this anti-inflammatory factor increased secretion. IL-8, also known as an anti-inflammatory protective factor, plays a role in promoting the migration of cells toward the injured area.

### UC-MSC secretome may have been applicable in wound healing of vascular abnormalities

Interestingly, our scratch test demonstrated that the Secretome derived from UC-MSCs could induce a chemotactic reaction in endothelial cells, increasing cell migration to the split site. This effect was observed in a time-dependent manner rather than being dependent on the concentration of UC-MSC secretome. A previous study by Caseiro and his colleague showed that the Secretome of UC-MSCs could stimulate migration in human endothelial cells in a scratch test (Caseiro *et al*., 2019). The chemotactic activity of the UC-MSC secretome is attributed to paracrine factors such as SDF-1, HGF, IL-8, and MCP-1, all of which contribute to migration and angiogenesis (Shen *et al*., 2015). Considering that the hCMEC/D3 cells express the MCP-1 or CCL2 receptor (CCR2) on their surface, it is reasonable to suggest that the enhancement of wound healing properties in the scratched area was mediated through this mechanism (Lopez-Ramirez *et al*., 2013)(Weber *et al*., 1999). Our recent research has shown that the treatment of hCMEC/D3 cells with αSN-AGs resulted in impaired collective cell movement (Hourfar *et al*., 2023). In this study we observed that the Secretome of UC-MSCs facilitated the migration of hCMEC/D3 cells in the scratch space, displaying a collective cell movement pattern, which is a notable result.

### αSN-AGs may directly lead to the initiation and spread of Parkinson’s disease by affecting on cerebral endothelial cells through blood side of brain

We propose some hypotheses to explain our observations on the protective role of Secretome against αSN-AGs. First, specific components of the Secretome might interact with αSN-AGs, preventing their attachment to the cell surface either through the protein corona or by interacting with the protein channels on the cell surface. Second, once inside the cells, the extracellular vesicles carrying proteins, lipids, or regulatory molecules (such as siRNA, miRNA, RNA, etc.) could protect by interfering with various cellular pathways involved in apoptosis, oxidative stress, inflammation, etc. Additionally, immune-modulating and anti-inflammatory factors secreted in the Secretome appear to moderate the system and reduce the intensity of the response. Accordingly, there is a possibility that the release of αSN-AGs from the blood to the brain can disrupt the BBB, even without the involvement of glial cells. This idea is supported by Braak’s proposal that Parkinson’s disease (PD) likely begins outside of the central nervous system and then spreads to the brain (Braak *et al*., 2003). In this respect, αSN-AGs have been shown to transfer from blood to the brain, and such transmission mainly occurs in Parkinson’s patients through exosomes (Sui *et al*., 2014).

Additionally, local bleeding in the brain of PD patients could initiate this damage and cause the αSN-AGs to spread to other parts of the brain. It is important to note that exosomes can easily pass through the BBB and enter the brain parenchyma through a process called endocytosis. Therefore, damage to the BBB is not solely a one-way process from the brain to the blood and does not exclusively involve glial cells (Matsumoto *et al*., 2017). However, the activation of glia and other cells within the NVU system in the later stages of damage expansion likely plays a significant role in determining the severity of the damage and the spread of symptoms.

Since the interaction between αSN-AGs and the BBB is reciprocal, and based on the findings of the current studies, we propose that if the BBB is compromised in a specific area of the brain, αSN-AGs can enter the bloodstream and potentially cause damage in another region of the brain using the previously mentioned mechanisms. Alongside the prion-like theory, this hypothesis provides an alternative perspective for understanding the spread of αSN-AGs in the brains of PD patients. Therefore, it is crucial to carefully consider the direct impact of αSN-AGs on the integrity of the BBB, particularly in the context of blood-to-brain transmission.

## Conclusion

In summary, the study’s results confirmed that αSN-AGs can directly cause dysfunction in the BBB without the need for other parts of the NVU to mediate.

Recent studies have shown that the Secretome of UC-MSCs can protect the BBB against damage induced by αSN-AGs, including mitochondrial Toxicity, apoptosis, nitrite oxide-induced toxicity, and dysfunctionality. Notably, the Secretome derived from UC-MSCs demonstrated greater efficacy in preserving the BBB’s integrity, measured by TEER and permeability, than the actual UC-MSCs in a BBB model. Moreover, it exhibited potent anti-inflammatory effects by releasing factors that combat inflammation and reducing the inflammatory factor TNF-α. These findings emphasize the growing preference for secretome-based cell-free therapy over traditional MSC-based cell therapies.

## Supporting information

Supplementary Results

## Acknowledgement

This study was supported by National Institute of Genetic Engineering and Biotechnology, Tehran, Iran (Grant No. 940701-I-523). D.E.O. is grateful for support from the Lundbeck Foundation. Denmark (Grant No. R276-2018671).

