## Supplementary Results for "Secretome of Human Umbilical cord mesenchymal stem cells exerts protective impacts on the blood-brain barrier against alpha-synuclein aggregates using an *in vitro* model"

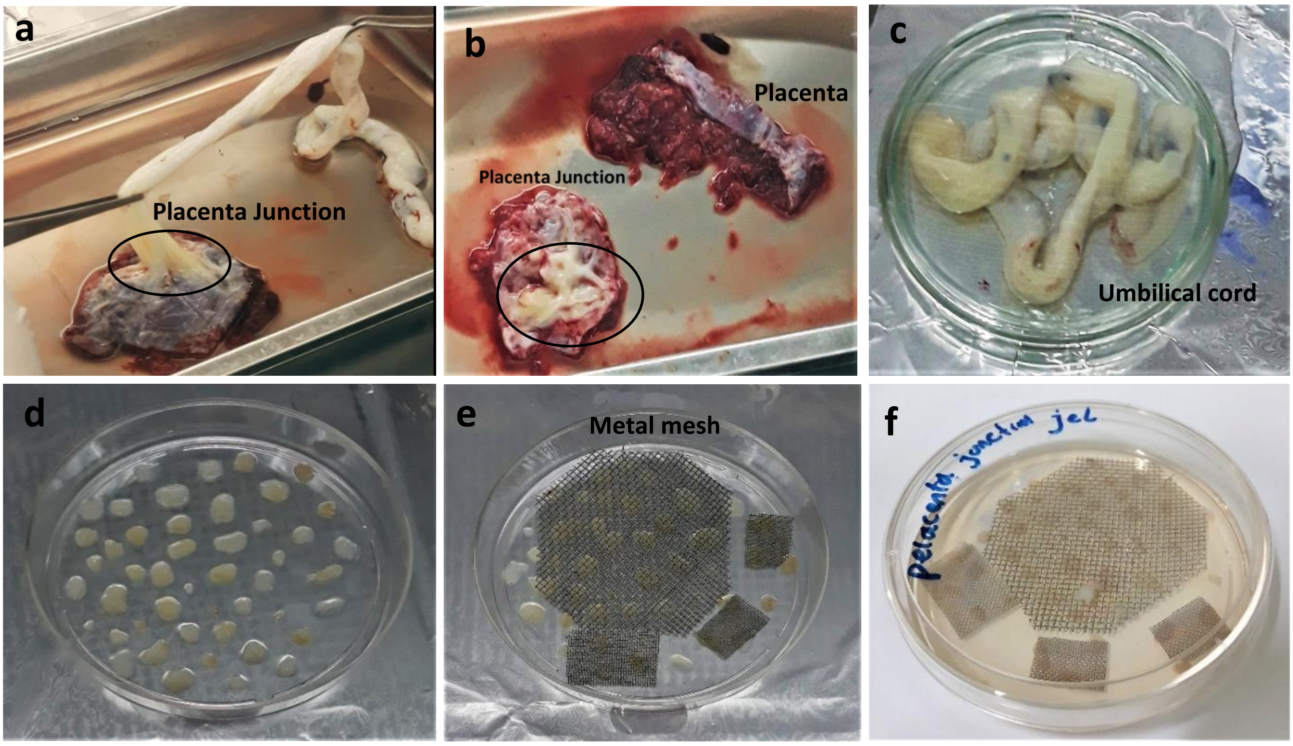


**Fig. S1**: **Procedure for MSCs isolation from umbilical cord tissue and placenta junction by the modified explant culture method.** (a) The circle marked part is the placenta junction, which is rich in MSCs. (b) The umbilical cord was separated from the end area connected to the placenta junction. Then the placenta area (red tissue below the placenta junction) and the placenta junction (circle marked part) were separated. (c) The separated umbilical cord was transferred into petri dish and divided into 1-2 cm pieces. (d) After washing and removing the vessels from the placental junction and the umbilical cord, each tissue was divided into 1-2 mm pieces and cultured on the bottom of the plates pre-coated with poly-L-lysine and gelatin. (e) Pre-sterilized metal meshes were placed on the cultured tissue pieces. (f) DMEM-F12 culture medium with 20% FBS and 2% antibiotics were added to the plates and transferred to a $CO_2$ incubator with 90% humidity.

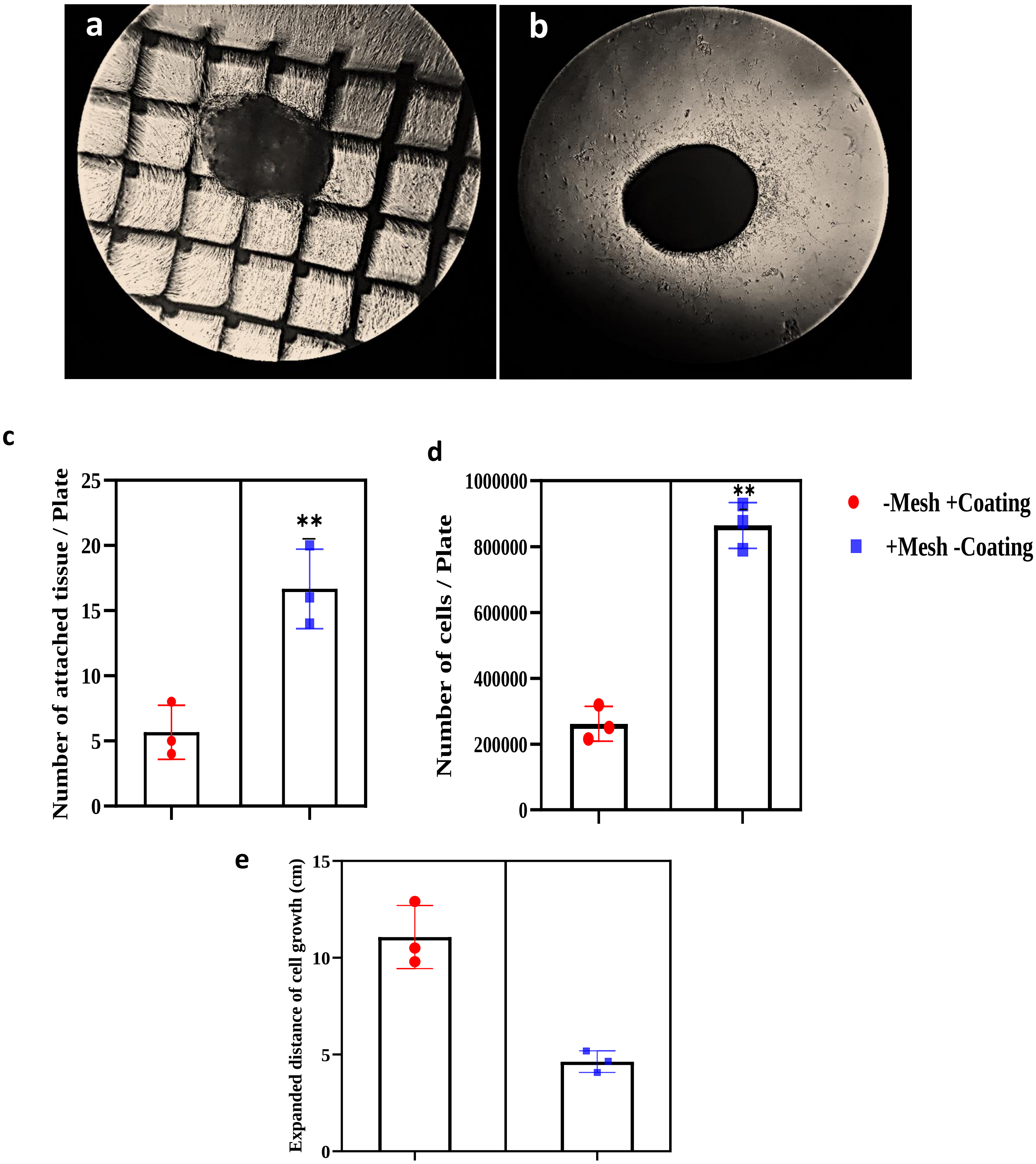


**Fig. S2**: Effect of some modifications in the procedure on the efficiency of MSCs obtained from the umbilical cord and placenta junction pieces. Comparing the amount of cell migration from the fixed tissues in the explant culture method (a) with or (b) without metal around the tissue is significantly high. Migrated mesenchymal cells with spindle-shaped morphology can be seen around the tissue with mesh, while without mesh, the number of cells around the tissue is deficient. (c) Assessing the number of tissue fragments attached to the bottom of the plates which release high migrated cells. The use of coating agent (poly-L-lysine and gelatin) on the bottom of the plates and the absence of metal mesh (red circle), and the use of metal mesh coating in the absence of coating (blue square) are shown. The initial number of cultured tissues in both plates was 20 pieces. (d) and (e) A comparison was made for the number of cells migrated and the area of moving (in cm). Metal mesh increases tissue attachment and quantity of the cell migration from tissue pieces. It has no impact on cell spread around each tissue piece. Coating the plate bottom prompts cell migration and growth in distant areas from each piece. The experiments were conducted thrice and analyzed using Digimizer software.

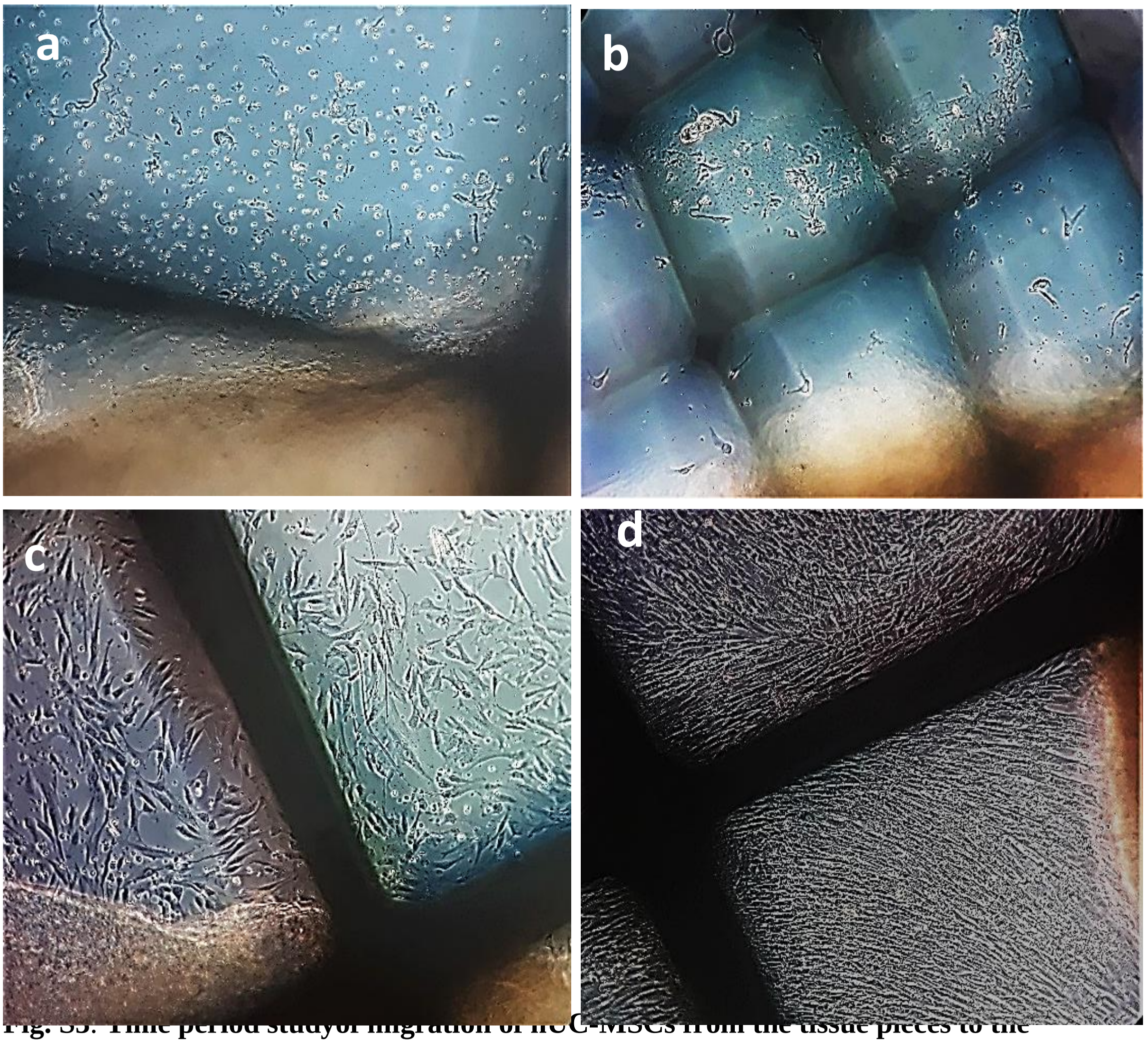


**Fig. S[illegible]. Time period studyof migration of hUC-MSCs from the tissue pieces to the surrounding area using the modified explant culture technique.** (a) On the 2nd to 5th day after tissue culture, cells with round morphology were observed around the tissues on the bottom of the coated plates along with the metal mesh. (b) On the 6th day, a sparse distribution of cells with characteristic morphology was identified. (c) On the 7th to 10th day after cultivation, the number of migrated cells with a spindle-shaped morphology increased rapidly. (d) On the 14th to 18th day after cultivation, the cells that migrated from the piece of tissue reached a suitable confluence for passage.

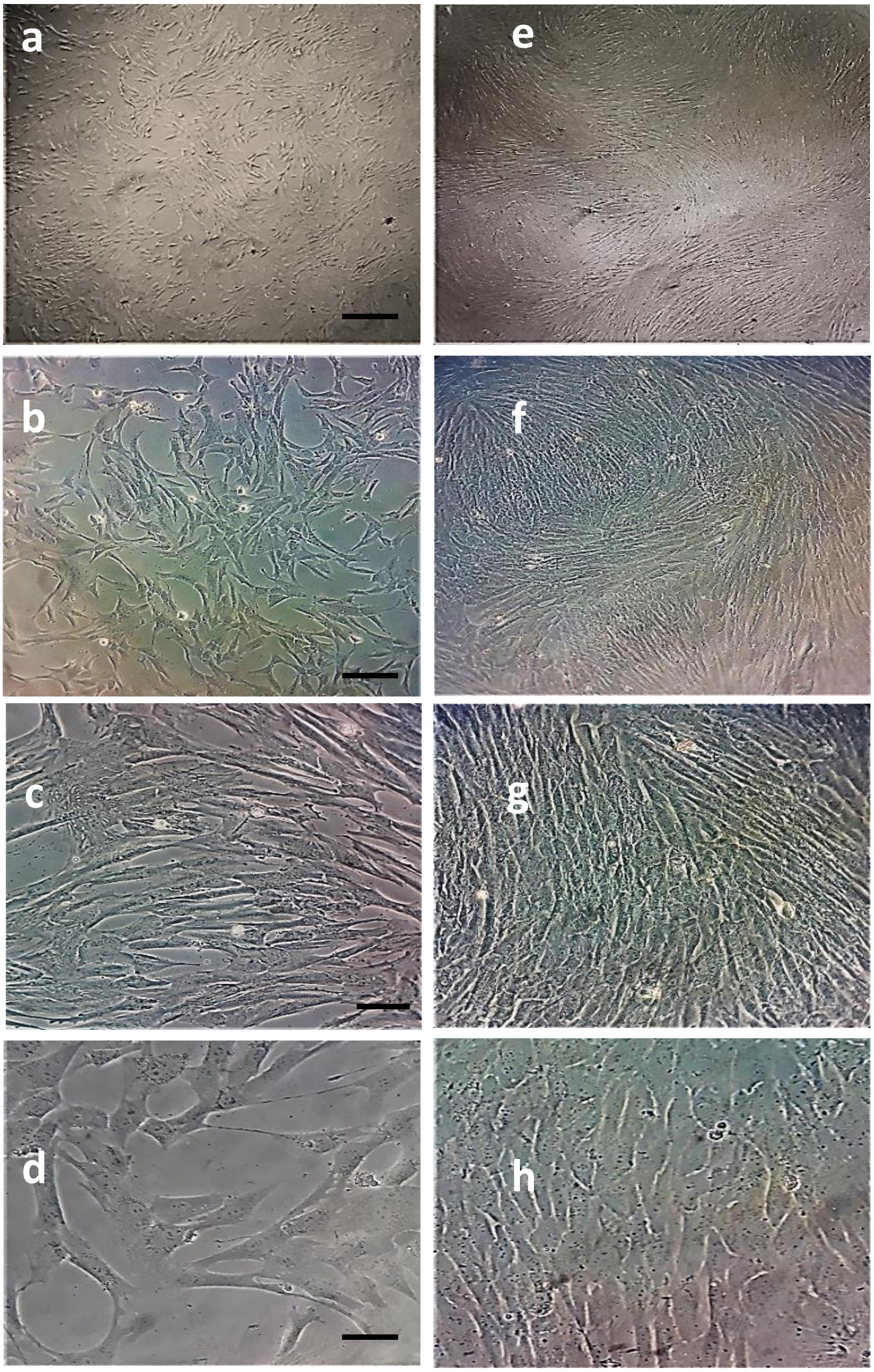


**Fig. S4**: **Evaluation of the morphology of MSCs extracted from the umbilical cord by phase contrast microscopy.** (a-d) The first passage, (e-h) the third passage before trypsinization and performing the flow cytometry assay. The morphology of the cells is spindle-shaped and fibroblast-like. Scale bar: 200 micrometers, lens magnification (a and e: 4X), (b and f: 10X), (c and g: 20X) (d and h: 40X).

a

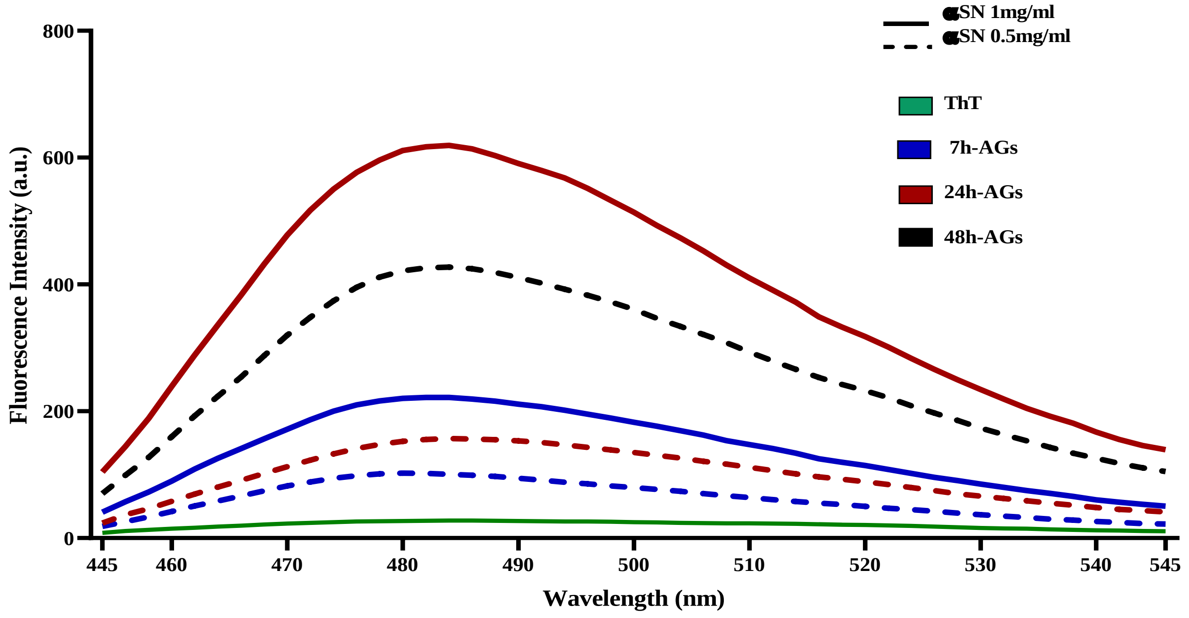
SN 1mg/ml
SN 0.5mg/ml
ThT
7h-AGs
24h-AGs
48h-AGs
800
600
400
200
0
Fluorescence Intensity (a.u.)
445
460
470
480
490
500
510
520
530
540
545
Wavelength (nm)

b

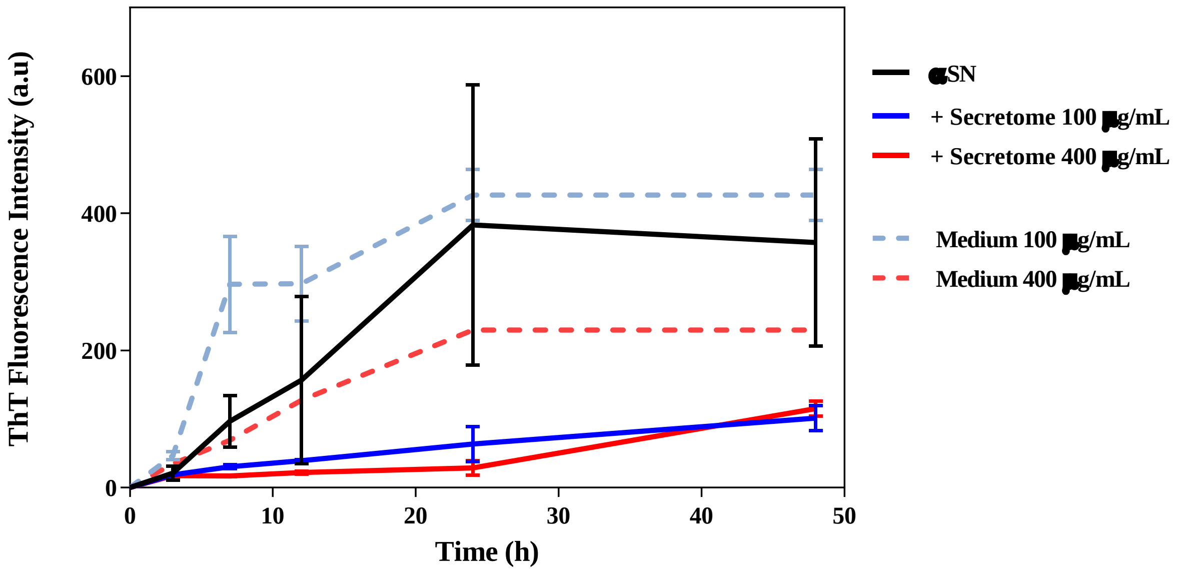
SN
+ Secretome 100 g/mL
+ Secretome 400 g/mL
Medium 100 g/mL
Medium 400 g/mL
600
400
200
0
ThT Fluorescence Intensity (a.u)
0
10
20
30
40
50
Time (h)

**Fig. S5**: **Exploring the αSN fibrillation and effect of the MSCs-derived secretome on the process.** (a) ThT fluorescence intensity of incubated αSN at two concentrations of 1 mg/mL (70 µM) and 0.5 mg/mL (35 µM) for 7, 24, and 48 hours. (b) Kinetic of the αSN fibrillation (1 mg/mL) in the presence and absence of the MSCs-derived secretome at two concentrations of 100 and 400 µg/mL. DMEM: RPMI culture medium in a ratio of 80:20 was also incubated with αSN during fibrillation. The MSCs-derived secretome at both 100 and 400 µg/mL concentrations inhibited αSN fibrillation.

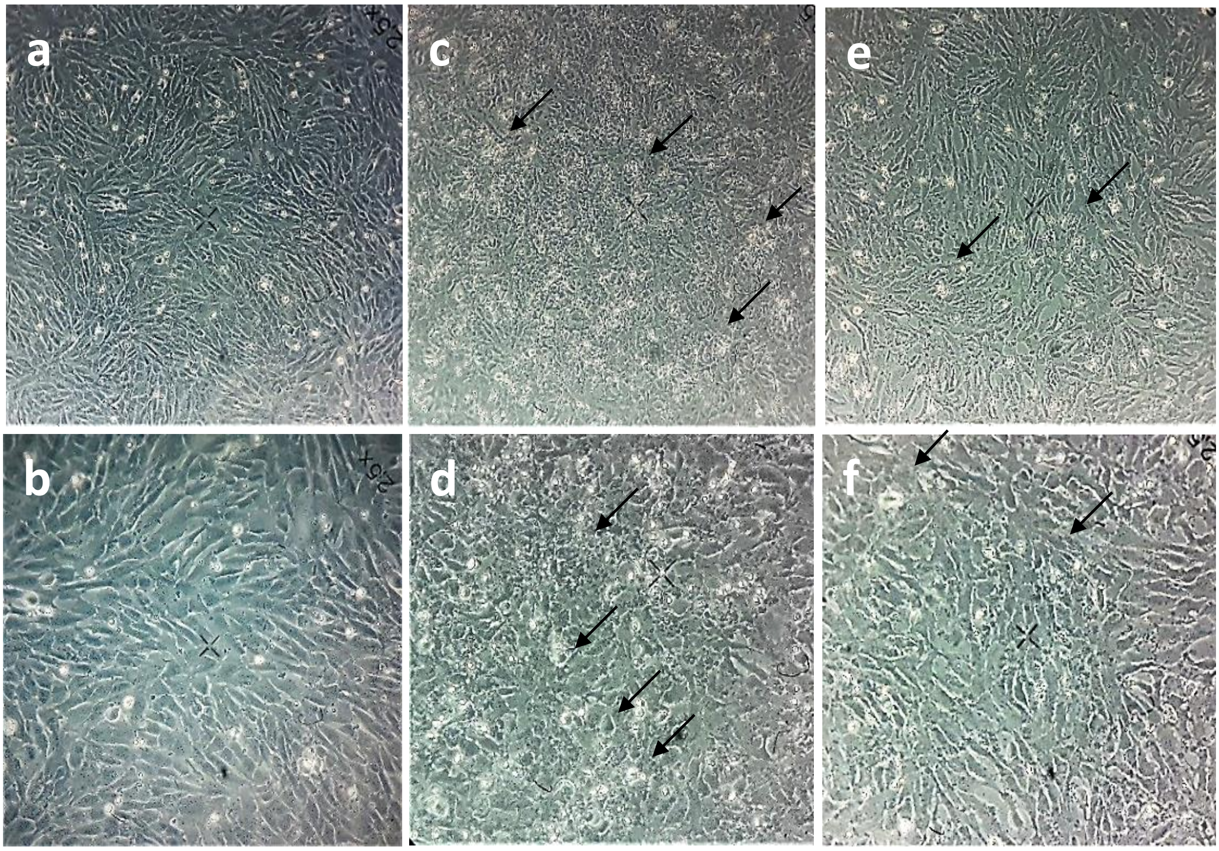


**Fig. S6**: **The effect of αSN-AGs in the absence or the presence of hUC-MSCs-derived secretome on the morphology of hCMEC/D3 cells.** (a) and (b) are control group before treatment; (c) and (d) are the hCMEC/D3 cells treatment with αSN-AGs (15% (v/v)) for 24 hours. Black arrows indicate some apoptotic cells. (e) and (f) are hCMEC/D3 cells treatment with αSN-AGs in the presence of hUC-MSCs-derived secretome with a volume ratio of 35% after 24 hours. The apoptosis of cells in the presence of secretome is significantly reduced (see fig 3j-v).

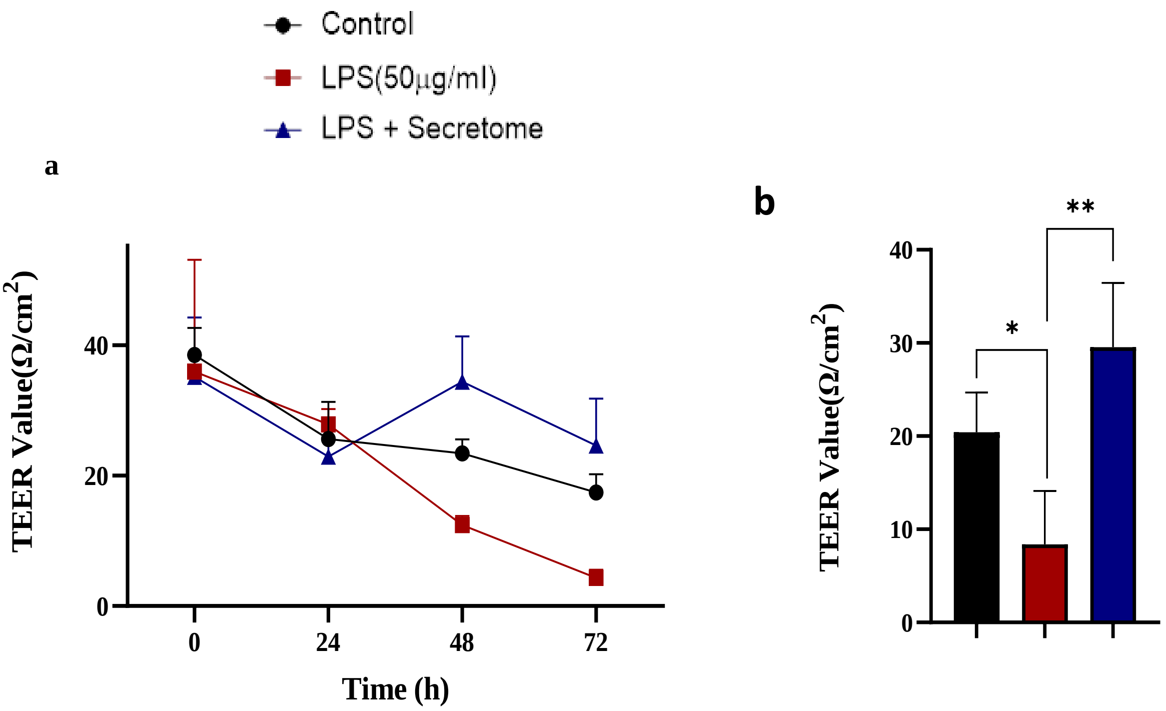


**Fig. S7**: **TEER measurement of the BBB monolayer model by culturing hCMEC/D3 cells after treatment with LPS in the absence and the presence of hUC-MSCs-derived secretome.** LPS with a concentration of 50 µg/mL was applied after the complete formation of a single layer of hCMEC/D3 cells on the apical region of the insert (after 7 days), and every 24 hours, TEER values were recorded with EVOM2 instrument (*P<0.05, **P<0.01). The presence of hUC-MSCs drive secretome at a concentration of 35% (v/v) significantly prevented the reduction of TEER caused by LPS toxicity. (a) TEER value measuring during the days, and (b) the graph is the final TEER obtained after 72 hours of treatment

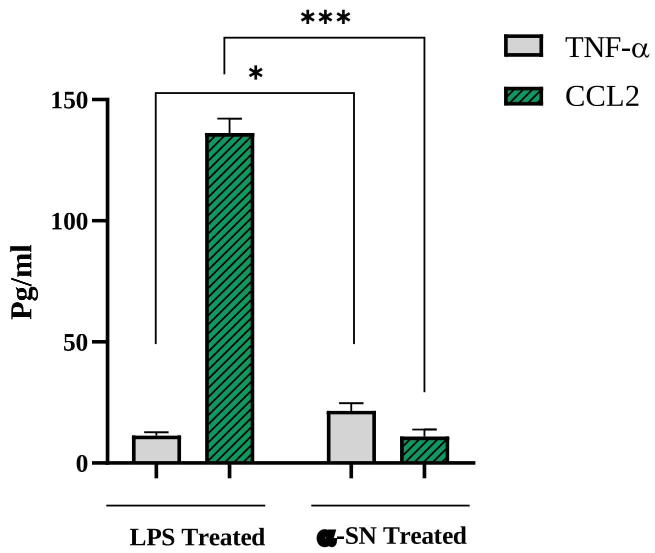

**Fig.S8**: **Investigation and comparison of the release of inflammatory factors, TNF-α and CCL2, from the BBB co-culture model after 13 hours of treatment with LPS and αSN-AGs using ELISA method.** The amount of CCL2 secretion triggered by LPS was considerably greater than that of TNF-α. Additionally, when αSN was administered after 24 hours, the release of TNF-α increased progressively over time. This indicates that LPS and αSN-AGs activate distinct inflammatory pathways.

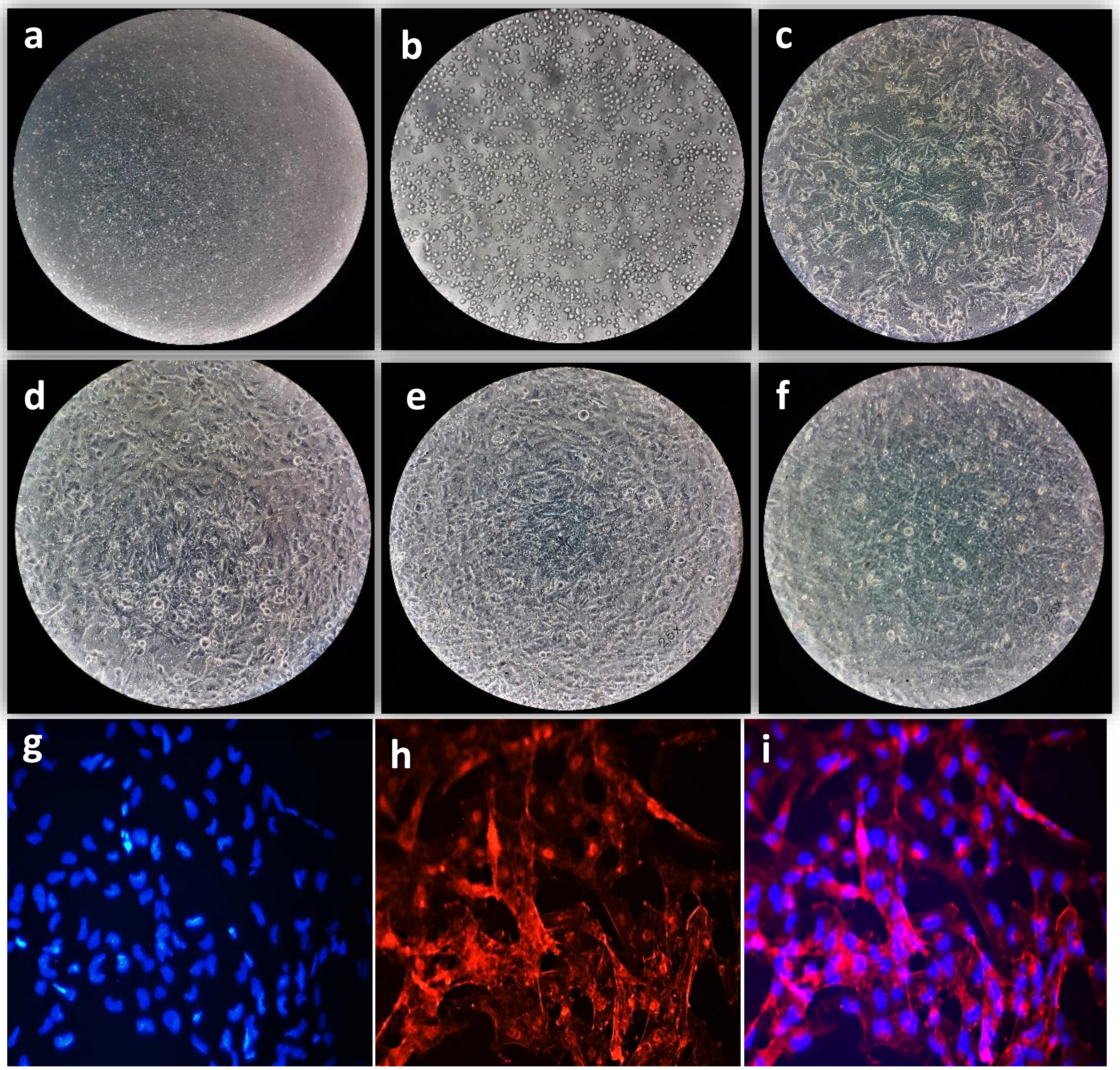

j

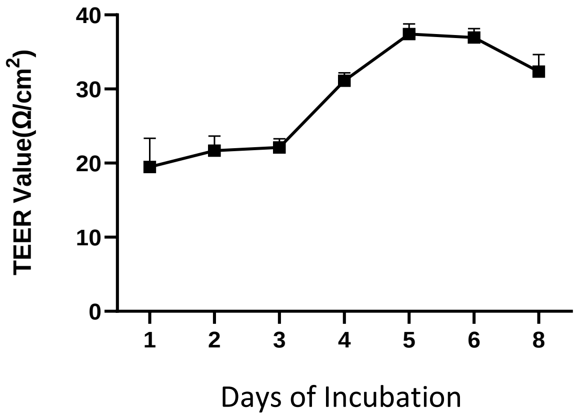


**Fig. S9**: **The stages of cell monolayer formation of the BBB model on the transwell insert**. (a) Transwell insert membrane with 0.4 µm pore size before coating with collagen. (b) Morphology of hCMEC/D3 cells in the first hours after cultivation. (c) The morphology of hCMEC/D3 cells after 24 hours of cultivation. d) Increasing the cell population after 48 hours of cultivation. e) Initiation of cell monolayer formation after 72 hours. (f) Complete formation of cell monolayer after 5 days. (g) and (i) hCMEC/D3 cells were stained with DAPI and Phaloidin dyes 24 hours after being cultured on a transwell insert. (g) Fluorescent microscope image of cells after staining with DAPI. (h) Fluorescent microscope image of the cytoskeleton of hCMEC/D3 cells after staining with Phaloidin. (i) Merge two images. (j) To determine the precise moment of cell monolayer formation, the TEER values were assessed daily for 8 days. During the third to the fifth day, there was an increase in TEER value, which can be attributed to the development of a cohesive cell layer. Up to sixth day, the TEER level remained nearly constant, indicating the successful establishment of a complete single cell layer resembling BBB model. However, from the sixth day onwards, a gradual decline in TEER level was observed. This suggests that the cells were transitioning from the single-layer state.

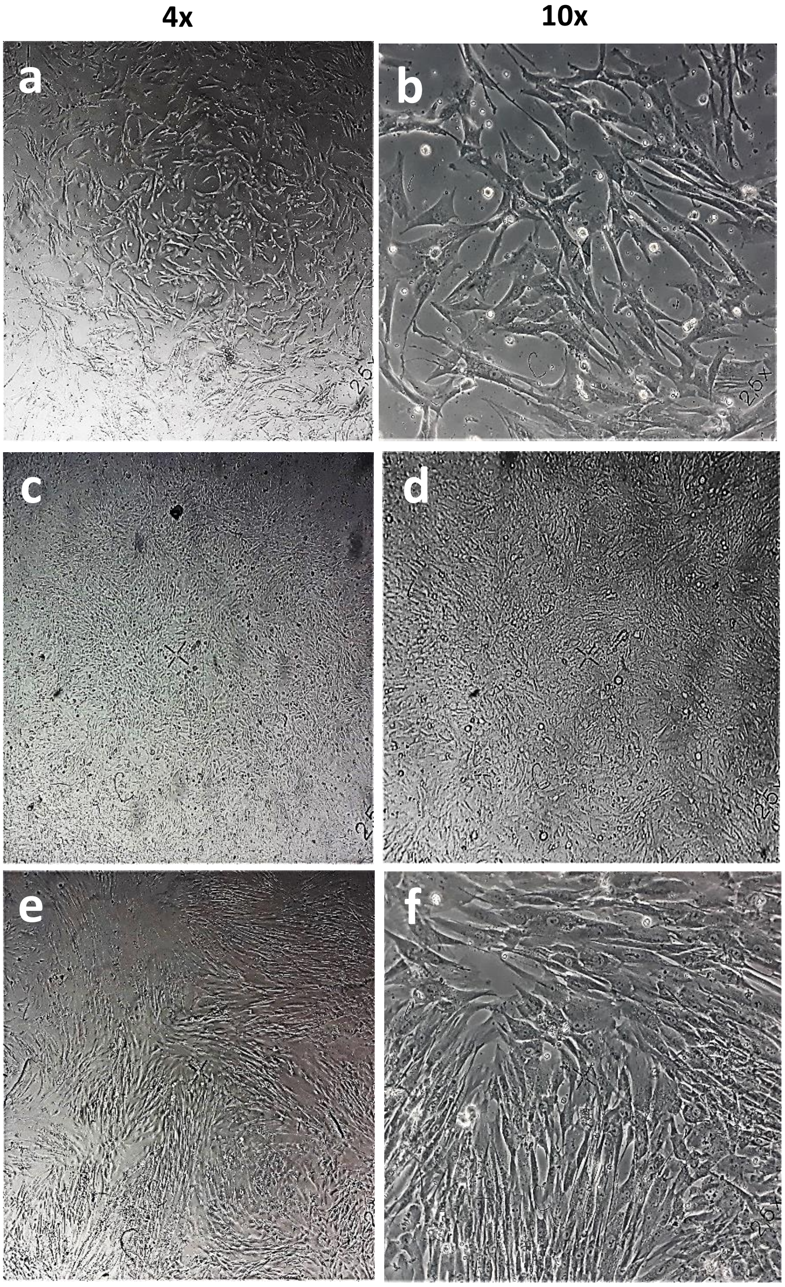


**Fig. S10**: **The morphology of hCMEC/D3 and UC-MSCs in a co-culture model of BBB.** (a) and (b) UC-MSCs were cultured one day earlier than hCMEC/D3 cells on the bottom of a 24-well plate. Morphology of UC-MSCs cells 24 hours after cultivation in DMEM culture media supplemented with 10% FBS. (c) and (d) Morphology of hCMEC/D3 cells, 48 hours after culture on transwell insert pre-coated with 0.1% collagen. (e) and (f) Morphology of UC-MSC cells before co-culture model treatment with αSN-AGs.
